# Structure of the complete dimeric human GDAP1 core domain provides insights into ligand binding and clustering of disease mutations

**DOI:** 10.1101/2020.11.13.381293

**Authors:** Giang Thi Tuyet Nguyen, Aleksi Sutinen, Arne Raasakka, Gopinath Muruganandam, Remy Loris, Petri Kursula

**Affiliations:** Faculty of Biochemistry and Molecular Medicine & Biocenter Oulu, University of Oulu, Finland; Department of Biomedicine, University of Bergen, Norway; VIB-VUB Center for Structural Biology, Vlaams Instituut voor Biotechnologie, Brussels, Belgium; Structural Biology Brussels, Department of Bioengineering Sciences, Vrije Universiteit Brussel, Brussels, Belgium

## Abstract

Charcot-Marie-Tooth disease (CMT) is one of the most common inherited neurological disorders. Despite the common involvement of ganglioside-induced differentiation-associated protein 1 (GDAP1) in CMT, the protein structure and function, as well as the pathogenic mechanisms, remain unclear. We determined the crystal structure of the complete human GDAP1 core domain, which shows a novel mode of dimerization within the glutathione S-transferase (GST) family. The long GDAP1-specific insertion forms an extended helix and a flexible loop. GDAP1 is catalytically inactive towards classical GST substrates. Through metabolite screening, we identified a ligand for GDAP1, the fatty acid hexadecanedioic acid, which is relevant for mitochondrial membrane permeability and Ca^2+^ homeostasis. The fatty acid binds to a pocket next to a CMT-linked residue cluster, increases protein stability, and induces changes in protein conformation and oligomerization. The closest homologue of GDAP1, GDAP1L1, is monomeric in its full-length form. Our results highlight the uniqueness of GDAP1 within the GST family and point towards allosteric mechanisms in regulating GDAP1 oligomeric state and function.

## INTRODUCTION

Mutations in the *GDAP1* gene, coding for the ganglioside-induced differentiation-associated protein 1 (GDAP1), are associated with several forms of Charcot-Marie-Tooth disease (CMT), which is one of the most common inherited neurological disorders, affecting 1 in 2500 people ^1–3^. GDAP1, a 358-amino-acid outer mitochondrial membrane (MOM) protein regulating the mitochondrial network, is highly expressed in neurons and less in Schwann cells ^4, 5^. GDAP1 contains two domains similar to the N- and C-terminal domains of glutathione (GSH) S-transferases (GST) (GST-N and GST-C, respectively), a hydrophobic domain (HD), and a transmembrane domain (TMD) ^6^. GDAP1 only shares ~20% sequence identity with canonical GSTs. Several GDAP1 constructs were previously assayed against a group of GST substrates, but no GSH-dependent activity or binding to GSH was detected ^7, 8^. However, a previous study suggested that GDAP1 has theta-class-like GST activity *in vitro*, which is regulated by the HD in an autoinhibitory manner ^6^.

Purified GDAP1 overexpressed in bacteria and insect cells forms dimers in solution, as shown by glutaraldehyde crosslinking and size-exclusion chromatography (SEC) ^6, 8^. Endogenous GDAP1 of human neuroblastoma SHSY5Y cells was detected in both dimeric and monomeric forms ^5^. The GDAP1 dimer disappeared under reducing conditions, implying that dimerization would be mediated *via* disulfide bonds. Contrary to these observations, the first crystal structure of the GDAP1 core domain, from a truncated construct lacking the large GDAP1-specific insertion, suggested that GDAP1 is monomeric ^7^. In light of these data, the GDAP1 insertion could play a role in GDAP1 dimerization and function.

GDAP1 functions in regulating the mitochondrial network by inducing fragmentation of mitochondria without inducing apoptosis. This fission activity is significantly reduced for CMT-related GDAP1 mutations ^4^. Recessive mutations in GDAP1 are associated with decreased fission activity, whereas dominant mutations induce impairment of mitochondrial fusion, increased production of reactive oxygen species (ROS), and susceptibility to apoptotic stimuli ^9^. To regulate various cellular processes, mitochondria use Ca^2+^ uptake and release to modulate cytosolic Ca^2+^ signalling ^10^. GDAP1 deficiency reduces Ca^2+^ inflow through store-operated Ca^2+^ entry (SOCE) activity and endoplasmic reticulum (ER)-Ca^2+^ levels ^11^. In the presence of Ca^2+^ or Sr^2+^, long-chain saturated α,ω-dioic acids, including hexadecanedioic acid (HA), can induce a cyclosporin A-insensitive permeability of the inner membrane of liver mitochondria ^12^.

The paralogous GDAP1-like protein 1 (GDAP1L1) shares a 55% sequence identity with GDAP1, and the HD and TMD are conserved ^13^. The HD and TMD are the targeting domains of GDAP1 for function in mitochondrial fission ^14^; however, GDAP1L1 is mainly cytosolic ^15^. GDAP1L1 can induce mitochondrial fission in the absence of GDAP1, implying that it may compensate for GDAP1 loss in the central nervous system ^15^.

Here, we describe the crystal structure of the complete dimeric GDAP1 GST core domain, and based on its unique mode of dimerization, we propose a model for full-length GDAP1 on the MOM. We also provide a low-resolution model for monomeric GDAP1L1 based on small-angle X-ray scattering (SAXS) data. As no GST activity was detected for GDAP1, a metabolite library was screened for GDAP1 binding partners. We find that HA binds to GDAP1 and affects its stability, conformation, and oligomerization. The HA binding site in the crystal structure of GDAP1 is located close to the CMT-linked residue cluster and the membrane-binding surface.

## MATERIALS AND METHODS

### Chemicals

Chemicals were from Sigma-Aldrich unless otherwise stated. Crystallization screens were from Molecular Dimensions. The Human Endogenous Metabolite Compound Library was from MedChemExpress.

### Cloning, expression, and purification

The open reading frame (ORF) of full-length human GDAP1 isoform 1 (UniProt ID: Q8TB36) was ordered from DNA 2.0 as a synthetic codon-optimized gene for bacterial cytosolic expression in the pJ201 vector. The C-terminally truncated GDAP1Δ319-358, GDAP1Δ303-358, and GDAP1Δ298-358 constructs were generated by PCR and transferred into the pDONR221 entry vector using Gateway^®^ technology-based site-specific recombination (Invitrogen). An N-terminal Tobacco Etch Virus (TEV) protease digestion site was included in each construct.

For structural and biochemical characterization, GDAP1 constructs were transferred into pTH27 ^16^ and pDEST-Trx ^17^ vectors, which encode for N-terminal His_6_ and thioredoxin tags, respectively. Point mutations were generated using site-directed mutagenesis PCR ^18^. The ORF of full-length GDAP1L1 was purchased in the pET28a(+)-TEV vector, containing a TEV protease cleavage site and a His_6_ tag (Genscript). All constructs were verified with DNA sequencing of both strands.

Recombinant protein expression was done using *E. coli* BL21(DE3) in ZYM-5052 auto-induction medium ^19^. Selenomethionine-substituted (SeMet) protein was expressed using *E. coli* B834(DE3) in SelenoMet^™^ -media (Molecular Dimensions) ^20^.

The soluble recombinant protein was captured on a Ni^2+^-NTA affinity resin by gravity flow (Thermo Fisher Scientific). Unbound proteins were washed with 25 mM HEPES, 300 mM NaCl, 2% glycerol, and 25 mM imidazole (pH 7.5). The protein was eluted with an identical buffer, with imidazole at 250 mM. The affinity tag was cleaved with a 1:20 molar ratio of TEV protease (16 h, +4 °C). The His_6_ -tag and TEV protease were then removed by another Ni^2+^-NTA affinity step. SEC was performed on a Superdex 200 or Superdex 75 10/300 GL increase column (GE Healthcare) using 25 mM HEPES (pH 7.5), 300 mM NaCl (SEC buffer) as eluent. An anion exchange chromatography (IEX) step was added for GDAP1L1, using a HiTrap HP Q XL column (GE Healthcare). GDAP1L1 was eluted using a linearly increasing gradient up to 1 M NaCl in 30 mM Tris (pH 7.9). Peak fractions were analyzed with SDS-PAGE, and Coomassie-stained bands were analyzed by Bruker UltrafleXtreme matrix-assisted laser desorption/time-of-flight mass spectrometer (MALDI TOF-MS). Tryptic peptides extracted from the gel were identified by searching NCBI and SwissProt databases using BioTools2.2 (Bruker).

### Crystallization, data collection, and structure determination

Crystallization was done using vapour diffusion at +4 °C. Protein and mother liquor drops were applied using a Mosquito LCP (TTP Labtech) nano-dispenser. The protein concentration was between 5-25 mg/ml in SEC buffer. Apo GDAP1Δ303-358 was crystallized in 0.2 M magnesium formate, 20% PEG 3350. GDAP1Δ303-358 crystals with HA were obtained by co-crystallization, with 1 mM HA (2% EtOH as solvent), and 0.1 M succinic acid, and 15% PEG3350 as mother liquor. SeMet-GDAP1 crystals were grown in 0.2 M ammonium formate, 20% PEG3350. Crystals were briefly soaked in 30% glycerol before flash-freezing in liquid N_2_. Data collection was conducted on the synchrotron beamlines P11 (DESY, Hamburg, Germany), I24, and I04 (Diamond Light Source, Didcot, UK), at 100 K (Table 4).

Data were processed and scaled with XDS ^21^ and AIMLESS ^22^. Phases were obtained from SeMet data with single-wavelength anomalous dispersion (SAD) using the Crank2 pipeline ^23^, and the initial model building was done using BUCCANEER ^24^. Molecular replacement, efinement, and structure validation were done using Phenix ^25, 26^ and CCP4 ^27^. The models were refined using Phenix.refine ^28^ or Refmac5 ^29^ and rebuilt using COOT ^30^. The GDAP1-HA complex structure was solved through molecular replacement with Phaser ^31^, using GDAP1 as a model. The structures were validated using MolProbity ^32^ and deposited at the PDB with entry codes 7ALM (apo) and 7AIA (HA complex).

### Bioinformatics and modelling

Structure visualization was done with PyMOL (http://www.pymol.org) and Chimera ^33^. Schematic views of the interactions were generated with LIGPLOT ^34^. Electrostatic surfaces were calculated with APBS and PDB2PQR ^35^. Structural homology searches were done using PSI-search ^36^ and SALAMI ^37^, and selected sequences were aligned with T-COFFEE ^38, 39^. Manual editing of the sequences was done using Genedoc and ESPRIPT3.0 ^40, 41^. Full-length GDAP1 modeling within a membrane was done using Coot and YASARA ^42^.

### Small-angle X-ray scattering

The GDAP1 monomer and dimer species were separated and analyzed with SEC-SAXS. Prior to the experiment, the samples were dialyzed against 25 mM HEPES pH 7.5, 300 mM NaCl and centrifuged at >20000 *g* for 10 min at +4 °C to remove aggregates. SAXS experiments were performed on the P12 beamline ^43^ (EMBL/DESY, Hamburg, Germany), the SWING beamline ^44^ (SOLEIL synchrotron, Saint Aubin, France), and the B21 beamline ^45^ (Diamond Light Source, Didcot, UK).

The data were collected over a s-range of 0.003–0.5◻Å^−1^ (s◻=◻*4*π *sin(*θ*)/*λ, where 2θ is the scattering angle) at a fixed temperature (+15 °C). 50 μl of a protein sample at 8.5-10 mg/ml was injected to a BioSEC3-300 (Agilent) or Superdex 75 10/300 GL increase column (GE Healthcare) and eluted a flow rate of 0.2◻ml/ml or 0.5 mg/ml. Data reduction to absolute units, frame averaging, and subtraction was performed using Foxtrot ^44^ or CHROMIXS ^46^.

Further processing and modeling were done using ATSAS 3.0 ^47^. Scattering curves were analyzed and particle dimensions determined using PRIMUS ^48^ and GNOM ^49^, and initial particle shape determination was performed using BODIES and AMBIMETER ^48, 50^. Chain-like models were generated using GASBOR ^51^. In combination with the crystal structure, hybrid modelling was performed using CORAL ^52^. CRYSOL ^53^ was used to evaluate fits of crystal structures to experimental data. SUPCOMB was used to superimpose SAXS models and crystal structures ^54^.

Low-resolution electron density reconstructions were calculated using DENSS ^55^. The electron density maps were calculated 20 times and averaged using EMAN2 ^56^. SAXS data and models were deposited at the SASBDB (Table S1).

### Multi-angle light scattering

Protein molecular mass and heterogeneity were determined by multi-angle light scattering (MALS) using a miniDAWN TREOS II detector (Wyatt Technologies), coupled to a Shimadzu Prominence HPLC system with RID-20A (RI) and SPD-M30A (diode array) detectors. SEC to separate oligomeric species was performed using Superdex 75 10/300GL or Superdex 200 15/150GL increase columns (GE Healthcare) in SEC buffer. The protein concentration was 1-10 mg/ml, and the injected protein amount 15-150 μg. Data processing, baseline reduction, and molecular weight calculation were done in ASTRA 7 (Wyatt Technologies).

### Thermal denaturation assays

GDAP1Δ319-358 and GDAP1Δ295-358 in SEC buffer were titrated with HA (final DMSO concentration 2% (v/v)) in a 96-well PCR plate. After adding SYPRO Orange fluorescent dye, the plate was sealed with an optical PCR plate sheet, and thermal denaturation was analyzed by differential scanning fluorimetry (DSF) in an Applied Biosystems 7500 device. Melting curves were analyzed with GraphPad Prism.

### Label-free stability assay

Thermal unfolding of wild-type and C88A GDAP1Δ303-358 in SEC buffer was studied by nanoDSF using a Prometheus NT.48 instrument (NanoTemper). The fluorescence of tryptophan was excited at 280◻nm and recorded at 330◻nm and 350◻nm. The samples were heated from +20 to +90 °C with a heating rate of 1 °C /min, and changes in the fluorescence ratio (F_350_/F_330_) were monitored to determine apparent melting temperatures (T_m_).

### Isothermal titration calorimetry (ITC)

The binding affinity of GDAP1Δ319-358 and GDAP1Δ295-358 towards HA was measured using a MicroCal iTC200 calorimeter (GE Healthcare) in SEC buffer with 2% (v/v) DMSO. The sample cell and injection syringe were filled with 50 μM GDAP1 and 500 μM HA, respectively. The system was equilibrated to a stable baseline before initiating an automated titration. The injection volume was 2.5 μl, and 15 injections were repeated at 180-s intervals at +25 °C. The sample was stirred at 750 rpm. The data were analyzed with the one-site binding model in Origin (MicroCal) to obtain thermodynamic parameters.

### Biolayer interferometry (BLI)

BLI measurements were performed in SEC buffer containing 0.005% Tween 20 and 2% DMSO, using an Octet RED instrument (FortéBio) at +25 °C. Biotinylated GDAP1Δ319-358 was loaded onto Super Streptavidin (SSA) biosensors (FortéBio) and quenched with 250 μl of 10 μg/ml biocytin. The association of GDAP1Δ319-358 with HA at a series of concentrations was measured for 180 s. The dissociation was performed by washing the biosensors with binding buffer for 180 s. A reference measurement without biotinylated protein was subtracted from all curves. Data were analyzed using Data Analysis 11.0 (FortéBio).

### GST activity assay

Spectrophotometric activity measurements were done using the generic GST substrate analogs 1-chloro-2,4-dinitrobenzene (CDNB), 4-nitrobenzyl chloride (pNBC), and 1,2-epoxy-3-(p-nitrophenoxy)propane (EPNP) together with every GDAP1 construct. Absorbance was followed at 360 nm for a 500-μl reaction at **+**25 °C for 5 min, with a Jasco V-730 UV-VIS spectrophotometer (JASCO International Co. Ltd., Tokyo, Japan) in a 1-mm quartz cuvette (Hellma Analytics). Substrate concentrations in the assays were 1 mM (CDNB), 0.25 mM (pNBC), and 0.3 mM (EPNP) in 100 mM potassium phosphate buffer, pH 6.5. The GSH concentration in was between 1-5 mM, and GDAP1 amount was 50 μg. As a positive control, 0.5 μg of recombinant *S. japonicum* GST-TEV fusion protein was used. Data were analyzed using Jasco Spectral analysis software. All measurements were done in triplicate.

## RESULTS

### Identification of a ligand affecting GDAP1 stability

We used the constructs GDAP1Δ295-358, GDAP1Δ303-358, and GDAP1Δ319-358 to get detailed insights into GDAP1 structure and potential functions (Figure 1A). Notably, the GDAP1-specific insertion (α-loop) was present in all constructs, in contrast to a recently reported GDAP1 structure ^7^.

**Figure 1.**
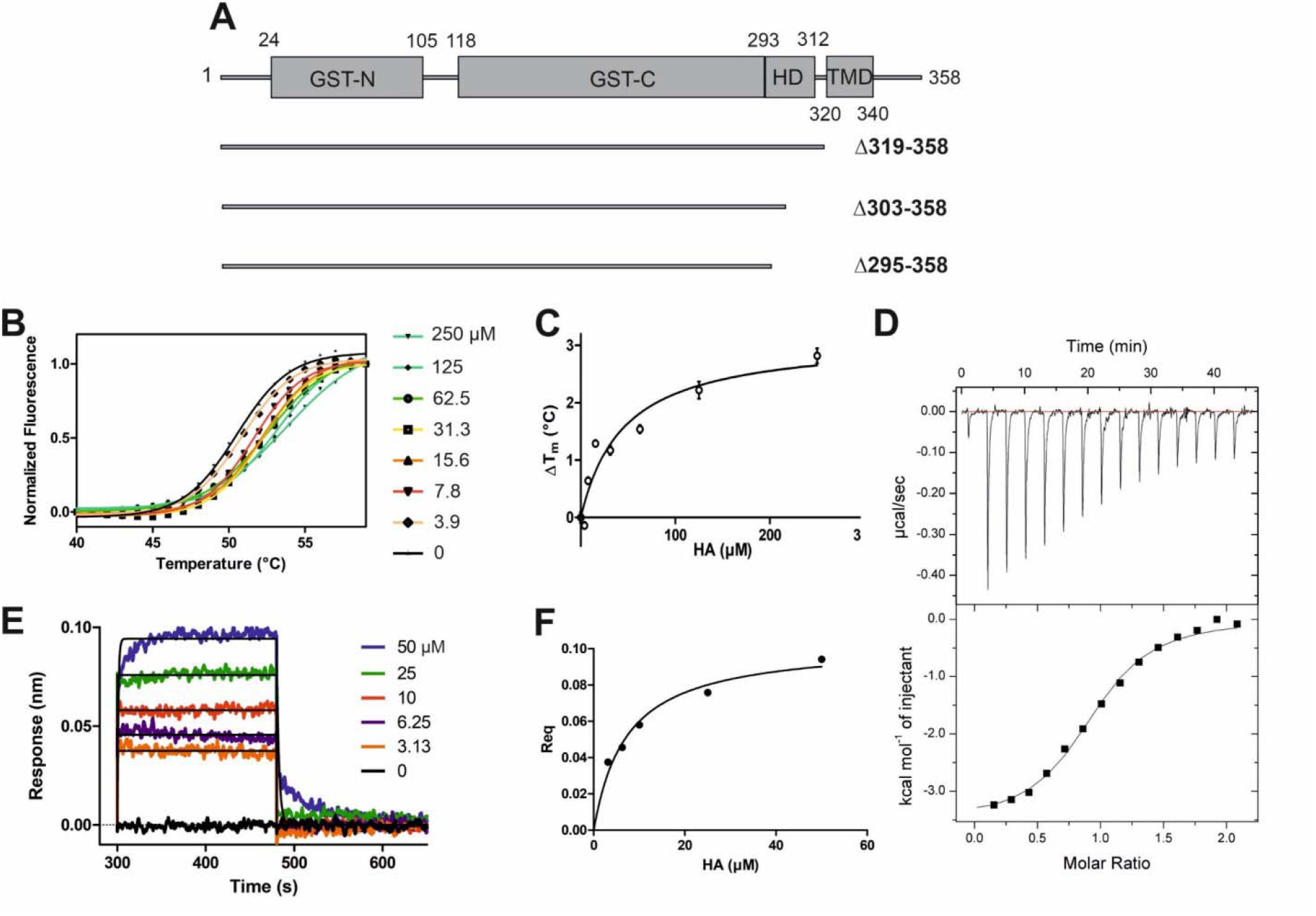
HA binding stabilizes GDAP1Δ319-358. A. Schematic of the full-length GDAP1 and three different deletion constructs used in this study. B. Thermal unfolding data. C. T_m_ shifts of GDAP1Δ319-358 upon HA titration. D. ITC binding curve of HA binding to GDAP1Δ319-358. E. BLI response of streptavidin-coated sensors derivatized with biotinylated GDAP1Δ319-358 and exposed to increasing concentrations of HA. F. Steady-state analysis of the BLI response versus HA concentration.

To search for ligands of GDAP1, compound library screening was performed using GDAP1Δ319-358 and GDAP1Δ295-358. Among ~300 compounds in a metabolite library, hexadecanedioic acid showed an effect on GDAP1 stability, increasing its T_m_ by ~3°C. Due to its limited solubility, HA was titrated up to 250 μM, and a concentration-dependent T_m_ shift was observed (Figure 1B, C, and S1A,B,C). The binding affinity of HA to GDAP1Δ295-358 and GDAP1Δ319-358 was determined using ITC and BLI (Figure 1D, E, F, and S1D). The K_d_ values determined by ITC and BLI are in the same range, whereas a higher K_d_ is detected using DSF for GDAP1Δ319-358 (Figure 1, Table 1). This could be due to an indirect effect from the fluorescent dye. Taken together, DSF, ITC, and BLI all show that HA binds to the GDAP1 core domain and stabilizes its structure.

**Table 1.** Binding affinities of HA to GDAP1 using different methods.

| **K<sub>d</sub> (μM)** | **GDAP1Δ319-358** | **GDAP1Δ295-358** |
| --- | --- | --- |
| DSF | 45.4 ± 18.3 | 9.7 ± 3.0 |
| ITC | 3.7 ± 0.2 | 2.0 ± 0.8 |
| BLI | 7.2 ± 1.5 |  |

### GDAP1 forms dimers in solution and HA binding affects protein oligomerization

GDAP1Δ295-358 and GDAP1Δ303-358 were subjected to synchrotron SEC-SAXS to investigate their oligomeric states and their conformation. The separation between dimer and monomer peaks is best for GDAP1Δ295-358 (Figure 2A and S2A). The linear fit in the Guinier region, the Porod volume, and the distance distribution function indicate monodispersity in the dimer peak of both constructs and the monomer peak of GDAP1Δ295-358, but the small second peak of GDAP1Δ303-358 appears to be a mixture of dimer and monomer. Clear separation of the monomer and dimer peaks enabled detailed analyses (Table 2, Figure 2B, C and S2B, C, D), and throughout this study, 3D modelling was only carried out for monodisperse, well-separated peaks. Figure 2D shows a chain-like *ab initio* dimer model of GDAP1Δ303-358.

**Figure 2.**
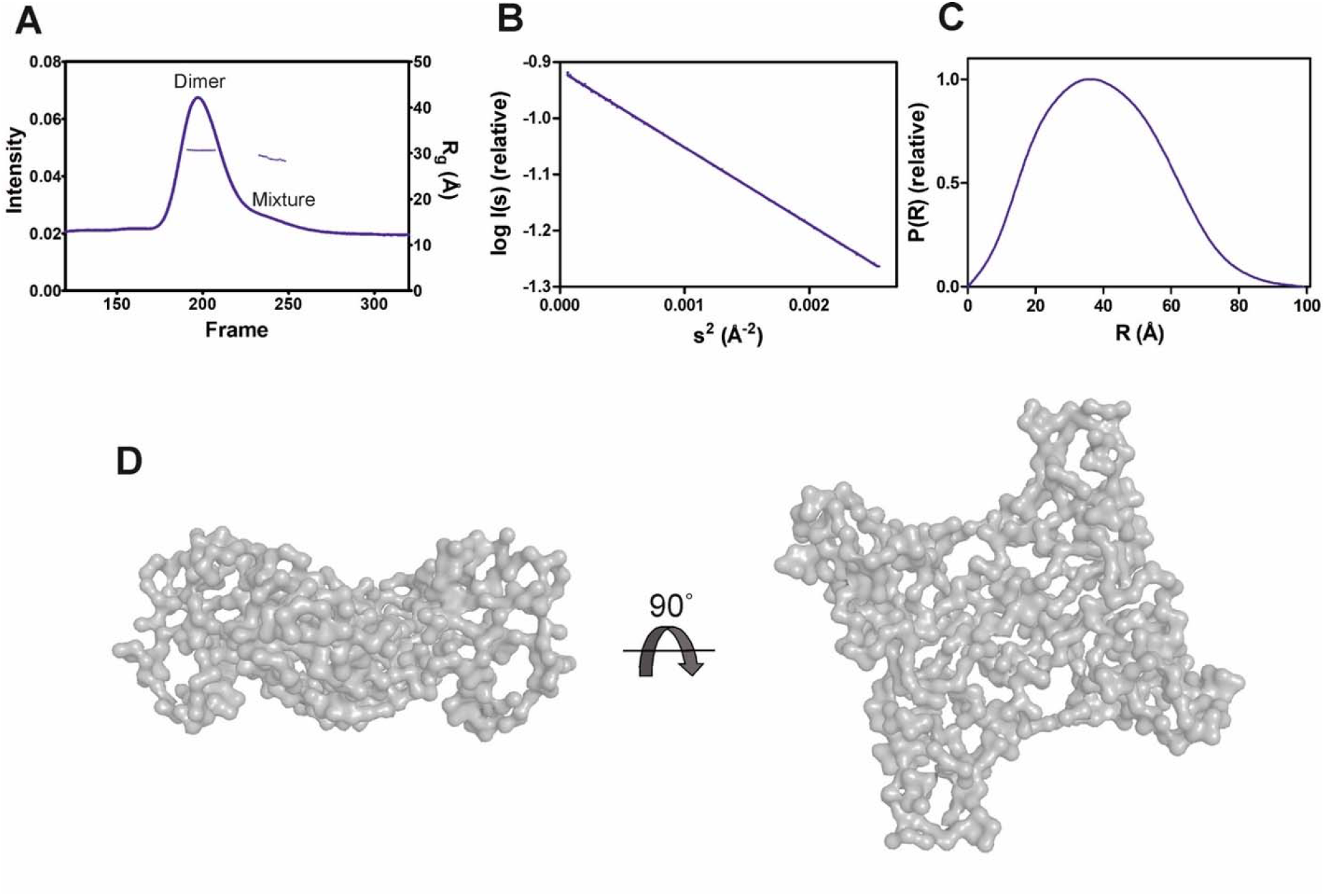
SAXS analysis of GDAP1Δ303-358. A. SEC-SAXS elution profile. R_g_ trace for the dimer and mixture of dimer/monomer peaks is also plotted. B. Guinier analysis of the dimer data. C. Distance distribution function for dimer. D. Two different views of the *ab initio* chain-like model of dimeric GDAP1Δ303-358.

**Table 2.**
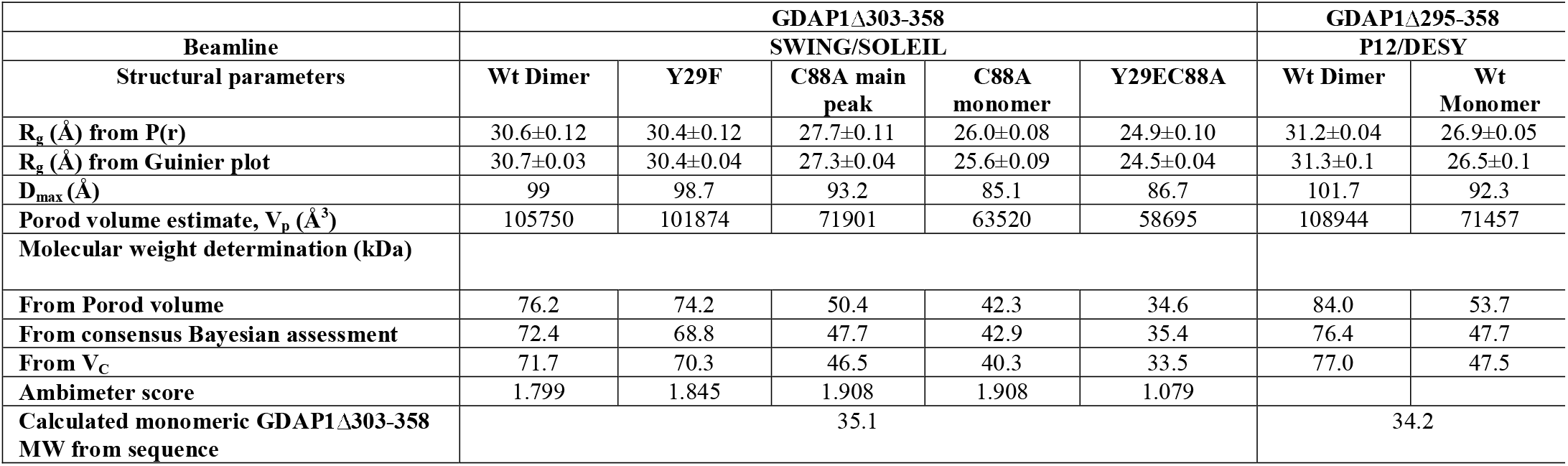
SAXS structural parameters of wild-type and mutant GDAP1Δ303-358 and GDAP1Δ295-358.

To assess particle shape ambiguity, the scattering curves were analyzed using AMBIMETER and BODIES without restraints (Table 2). The slight score value variation suggests that both species are likely homogeneous and monodisperse, which agrees with the distance distribution functions.

To examine the concentration dependence of GDAP1 oligomerization, we tested two different concentrations for each construct using SEC. At lower concentrations, SEC data show two peaks corresponding to dimers and monomers, whereas broad peaks are observed at higher concentrations, implying a dimer/monomer equilibrium (Figure S3). The main peak of GDAP1Δ319-358 is a mixture of dimer and monomer and could only be separated at a very low concentration (Figure S3B). Under non-reducing conditions, the protein adopts both dimeric and monomeric forms (Figure S3D). Dimers are not detected under reducing conditions (Figure S3D), indicating an inter-subunit disulfide bond involved in dimerization. These results are consistent with earlier observations that the endogenous GDAP1 dimer disappears in the presence of dithiothreitol (DTT) ^5^. Monomeric and dimeric GDAP1 can nevertheless be present in solution in a dynamic equilibrium as the dimer seems to form transiently and is dependent on the redox state.

To test the effect of HA on GDAP1 oligomerization, we performed SEC-SAXS using GDAP1Δ295-358 and GDAP1Δ319-358. The elution profile of the protein-HA complex shows a higher monomer fraction than apo GDAP1 (Figure 3A and S4A). The apo GDAP1Δ319-358 has a broad peak containing both dimer and monomer, whereas the complex elutes as two well-separated peaks (Figure 3A, Band S4A, B). Hence, HA binding allowed us to analyze monomeric and dimeric GDAP1 separately by SEC-SAXS.

**Figure 3.**
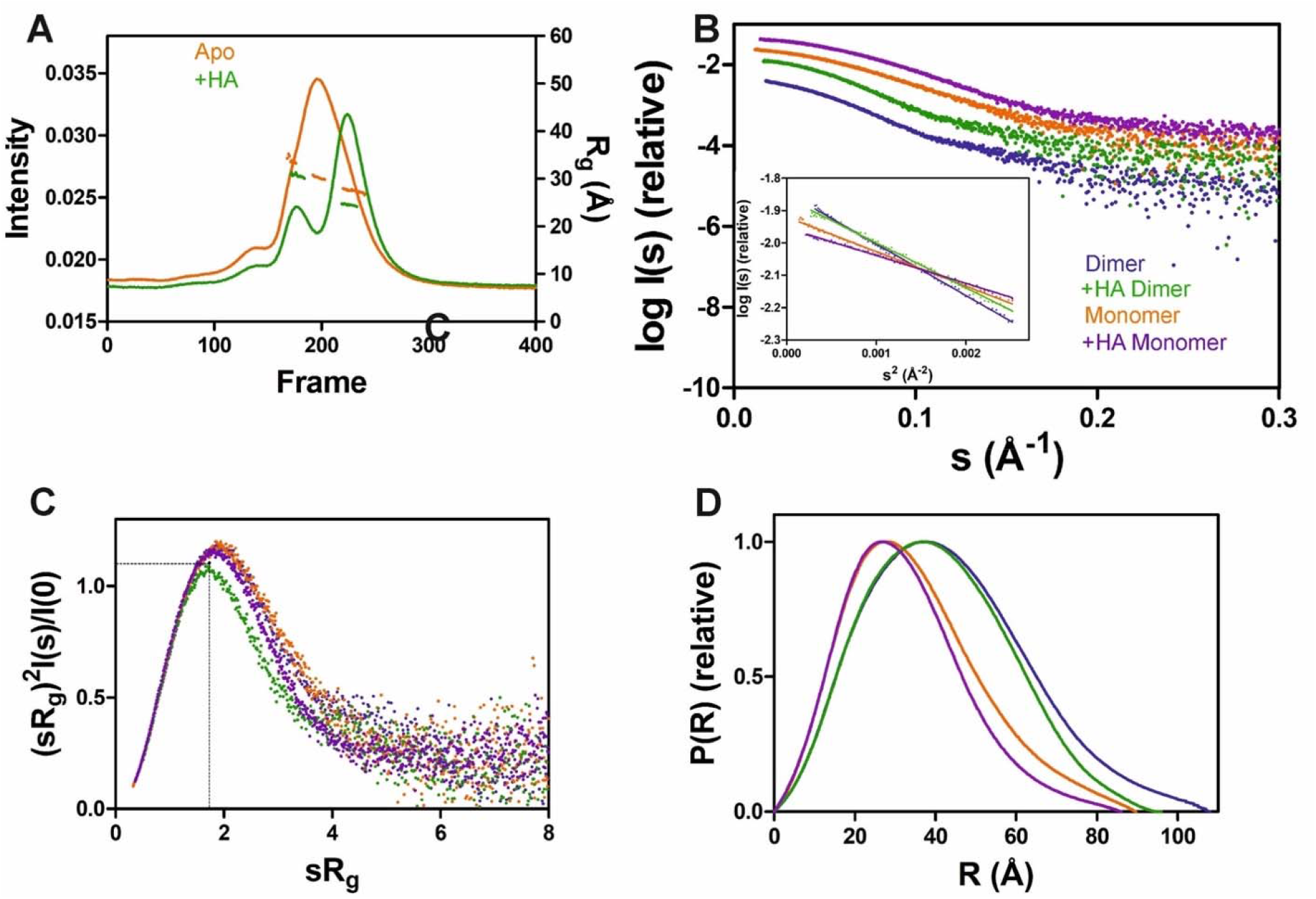
SAXS analysis of GDAP1Δ319-358 in the absence and presence of HA. A. SEC-SAXS elution profiles and R_g_ plot of SAXS frames for dimer and monomer peaks of the protein. B. Experimental scattering data (log (I_s_) vs. s) and Guinier analysis (inset). C. R_g_ normalized Kratky plots, the dashed lines representing the maximum value of a standard globular protein. D. Distance distributions *p(r)* plots of ligand-free GDAP1 dimer (blue) and monomer (orange), ligand-bound GDAP1 dimer (green) and monomer (purple).

In a dimensionless Kratky plot ^57, 58^, folded globular proteins show a bell-shaped curve reaching its apex of 1.1 when 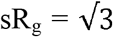, and multidomain proteins connected by linkers with a compact overall conformation have a bell-shaped curve, which is asymmetrically stretched ^59^. The higher the sR_g_ value at the apex of the curve, the greater the flexibility and disorder of the protein ^57^. Dimensionless Kratky plots suggest that apo GDAP1 is less compact than HA-bound GDAP1 (Figure 3C and S4C), and distance distributions, as well as R_g_, indicate compaction of GDAP1 upon ligand binding for both monomeric and dimeric GDAP1 (Figure 3D, S4D and Table 3). Hence, the stabilization of the GDAP1 structure is accompanied by a more compact 3D structure.

**Table 3.** SAXS structural parameters of GDAP1Δ319-358 and GDAP1Δ295-358 in the absence or presence of HA.

| Structural parameters | GDAP1Δ319-358 |  |  |  | GDAP1Δ295-358 |  |  |  |
| --- | --- | --- | --- | --- | --- | --- | --- | --- |
|  | SWING /SOLEIL |  |  |  |  |  |  |  |
|  | Apo<br>Dimer | Apo<br>Monomer | +HA<br>Dimer | +HA<br>Monomer | Apo<br>Dimer | Apo<br>Monomer | +HA<br>Dimer | +HA<br>Monomer |
| R<sub>g</sub> (Å) from P(r) | 33.5±0.01 | 27.3±0.01 | 31.2±0.01 | 25.1±0.01 | 32.2±0.01 | 29.3±0.01 | 30.9±0.01 | 26.1±0.01 |
| R<sub>g</sub> (Å) from Guinier plot | 34.7±0.22 | 27.1±0.08 | 30.9±0.14 | 24.6±0.05 | 32.4±0.07 | 29.1±0.09 | 30.7±0.08 | 25.3±0.09 |
| D<sub>max</sub> (Å) | 107.9 | 89.6 | 96 | 86 | 100.6 | 93.5 | 92.1 | 88.6 |
| Porod volume estimate, V<sub>p</sub> (Å<sup>3</sup>) | 129499 | 69390 | 120949 | 64543 | 117837 | 78347 | 112145 | 63005 |
| DAMMIN model volume | 142660 | 84395 | 136260 | 80269 | 132120 | 98337 | 124560 | 80269 |
| χ<sup>2</sup> against raw data for GASBOR models | 1.29 | 1.34 | 1.11 | 1.38 | 1.22 | 1.19 | 1.36 | 1.20 |
| Molecular weight determination (kDa) |  |  |  |  |  |  |  |  |
| From Porod volume | 97.6 | 46.3 | 85.3 | 36.9 | 83.7 | 57.8 | 77.6 | 40.2 |
| From DAMMIN model volume | 71.3 | 42.2 | 68.1 | 40.1 | 66.1 | 49.2 | 62.3 | 40.1 |
| From consensus Bayesian assessment | 94.2 | 46.7 | 85.6 | 39.4 | 80.8 | 58.2 | 74.3 | 40.2 |
| From V<sub>C</sub> | 89.9 | 44.4 | 81 | 37.5 | 77.3 | 53.5 | 73 | 40.8 |
| Calculated monomeric MW from sequence | 36.9 |  |  |  | 34.2 |  |  |  |

### Crystal structures of apo and liganded GDAP1 reveal structural relations to the GST family but suggest lack of GST activity

We determined the crystal structure of human GDAP1Δ303-358 at 2.8-Å resolution and its complex with HA at 2.2-Å resolution. Notably, this complete GDAP1 core domain, containing the GDAP-specific ⍰-loop insertion, assembles as a homodimer (Table 4, Figure 4A).

**Figure 4.**
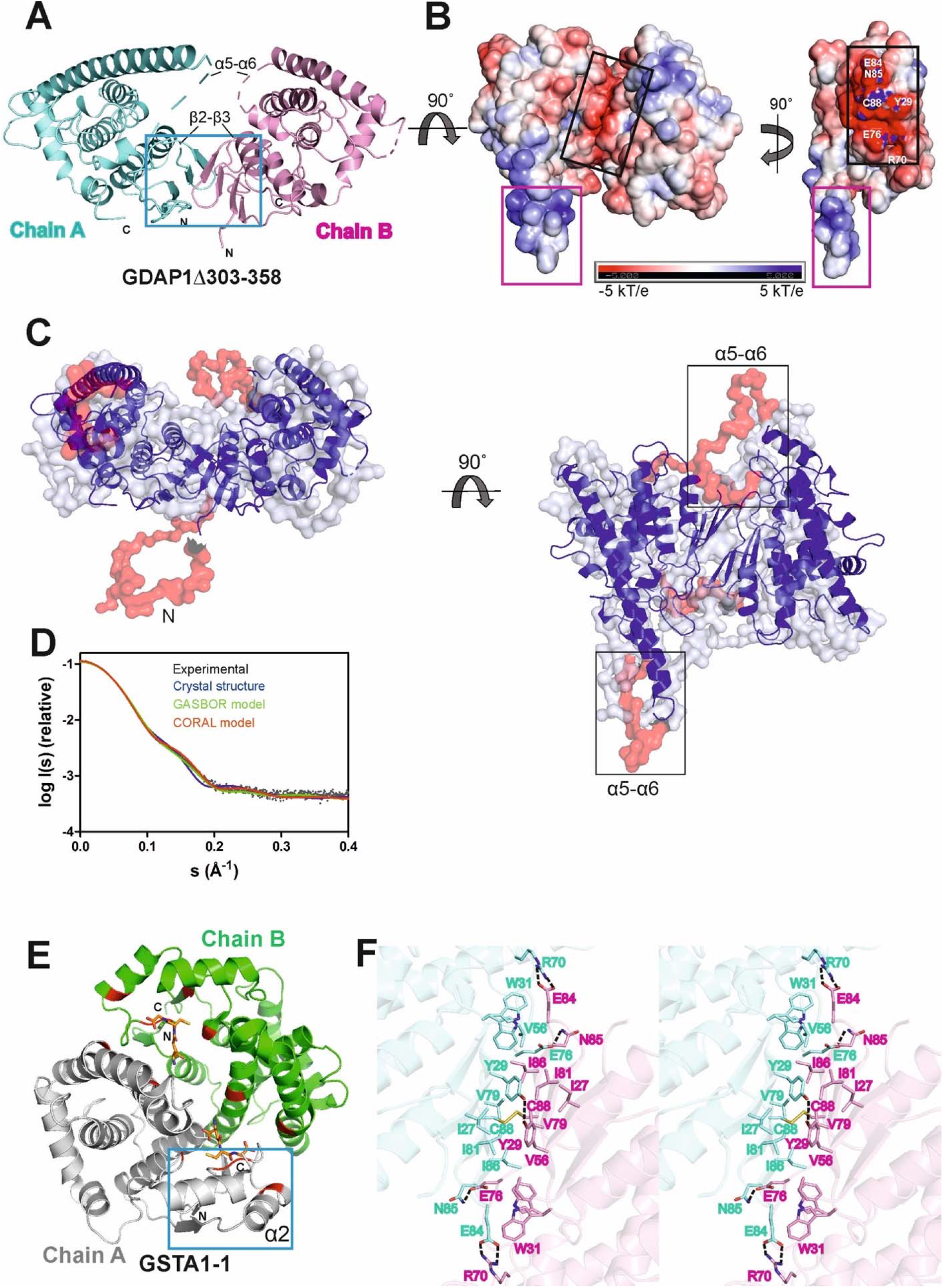
Crystal structure of the complete GDAP1 core domain and the dimer interface of GDAP1 compared to GSTA1-1. The orientation of GDAP1 chain A (cyan) and GSTA1-1 chain A (grey) are kept the same. A. Overall structure of the dimeric GDAP1Δ303-358. Chain A and chain B are shown in cyan and pink, respectively. The dashed lines indicate loops not defined by electron density. The dimerization interface is highlighted with a blue box. B. Electrostatic surface potential of GDAP1Δ303-358 is presented. The rotation shows only chain A. Note the strong positive potential of the long helix α6 (magenta box) and the negative potential of the dimer interface (black box). C. Dimeric GSTA1-1 ^65^. GSH is shown as orange sticks. The catalytic residues are shown in red. Blue box is the GST-N region corresponding to GDAP1 dimer interface. Chain A and chain B are shown in grey and green, respectively. D. Stereo view of the dimer interface showing interacting residues. The disulfide bond is shown, and hydrogen bonds are indicated by dashed lines. E. Two different views of the *ab initio* chain-like model of dimeric GDAP1Δ303-358 (transparent surface) superimposed with the GDAP1 crystal structure (blue) and the hybrid model. The built regions not present in the crystal structure are shown in red. F. Experimental scattering curve of GDAP1Δ303-358 dimer (black) overlaid with the theoretical scattering curve calculated from GDAP1Δ303-358 structure (blue, χ^2^ =17.1), GASBOR model (green, χ^2^ =38.2) and CORAL model (red, χ^2^ =4.7) using CRYSOL.

Similarly to other GST family members, each GDAP1 monomer includes an N-terminal thioredoxin-like domain and a C-terminal α-helical domain. The GST-N domain has four β strands, forming a β sheet and two α helices with the topology β1-α1-β2-β3-β4-α2 (Figure S5A, C), whereas, in canonical GST ^60^, an additional α helix between β2 and β3 is present, forming an overall topology β1-α1-β2-α2-β3-β4-α3 (Figure S5B, D). The GST-C domain is composed of seven α helices with a long α6 helix, visible in in one monomer of the dimer. In the other chain, this helix is shorter (Figure 4A, B), implying flexibility of the α6 helix and a local breakdown of non-crystallographic symmetry. The β2-β3 loop, residues Ser73-Val77, and the α5-α6 loop, Gln163-Glu183, do not display clear electron density, also indicating flexibility (Figure 4A) even though the α5-α6 loop region was predicted to contain an additional α helix ^7^. The electrostatic potential map reveals mainly a negative charge close to the dimer interface, whereas a strong positive charge is found on the exposed surface of the long helix α6 (Figure 4B).

The chain-like SAXS *ab initio* dimer model superimposes well with the crystal structure (Figure 4C). A hybrid model of GDAP1Δ303-358 was generated based on the crystal structure, building the missing residues (Figure 4C). This hybrid model fits the experimental SAXS data better than the chain-like model or a theoretical scattering curve generated from the crystal structure (Figure 4D). Hence, the conformation of GDAP1 in solution closely resembles that in the crystal state, and a simple rebuilding of the missing segments reproduces the solution scattering curve.

To complement the SAXS analysis, electron density reconstructions were prepared using DENSS ^55^ from GDAP1Δ303-358 SAXS data. According to the averaged DENSS electron density map (Fig. S6), the conformational difference between the two subunits of the dimer in the crystal seems to also exist in solution. The α5-α6 loop is visible in the map, supporting the rigid body model of the missing loops. Although the dimer interface is small, the dimer is stable in solution. The particle dimensions computed from the maps agree with the distance distribution functions (Table S2).

In contrast to canonical GST dimer interface contacts between GST-N of one subunit and GST-C of the other, involving β4, α3, α4, and α5 (Figure 4E), the dimer interface of GDAP1 forms entirely between the GST-N domains (Figure 4A, F). The interactions at the GDAP1 dimer interface include a disulfide bond between the Cys88 residues in strand β4 and a hydrogen bond between Tyr29 in strand β1 of each monomer (Figure 4F, S7A). Moreover, ion-dipole interactions between the Asn85 and Glu76 sidechains, as well as a salt bridge between Glu84 of one monomer and Arg70 of the other monomer, contribute to dimer formation (Figure 4F, S7A). Together with Tyr29, many residues, including Ile27, Tryp31, Val56, Val79, Ile81, and Ile86, create a hydrophobic surface at the dimer interface (Figure 4F, S7A). The above observations are consistent with the lack of GST activity in GDAP1, as the canonical dimerization mode generates the active site with sites for substrate binding. We could not detect any activity towards the conventional GST substrates CDNB, NBC, and EPNP, even at high protein concentration (Table S3). Similarly, previous publications showed no GSH binding to GDAP1 using ITC ^7^ or no GSH-dependent activity ^8^.

The crystal structure of GDAP1 in complex with HA (Table 4) reveals the ligand-binding site. HA binds to a pocket in the C-terminal domain formed by helices α1, α8, and α9 and their connecting loops (Figure 5A). The side chains of Arg282 and Gln235, together with Lys287 and Arg286, make hydrogen bonds and salt bridges to the carboxyl groups of HA, whereas the alkyl moiety forms van der Waals interactions with residues lining the pocket, including Trp238, Phe244, and Thr288 (Figure 5B, C, and Figure S7B). Superposition of our complex and apo structures with truncated mouse GDAP1 ^7^ and schistosomal GST ^60^ shows differences in loops β2-β3, α5-α6, and α6-α7 (Figure 5D). The loop β2-β3 of GDAP1 becomes more ordered in the presence of the ligand, whereas no electron density is present in this region in the apo structure and the mouse GDAP1. The loop β2-β3 contains the α2 helix in the canonical *Sj*GST structure (Figure 5D and S5). The α5-α6 loop in human GDAP1 is a unique structure compared to the truncated mouse GDAP1 and *Sj*GST (Figure 5D). The α6-α7 loop shift makes the structure more compact in the presence of the ligand (Figure 5D). This observation is consistent with SAXS data, implying more compact conformations in the presence of HA (Figure S8).

**Table 4.**
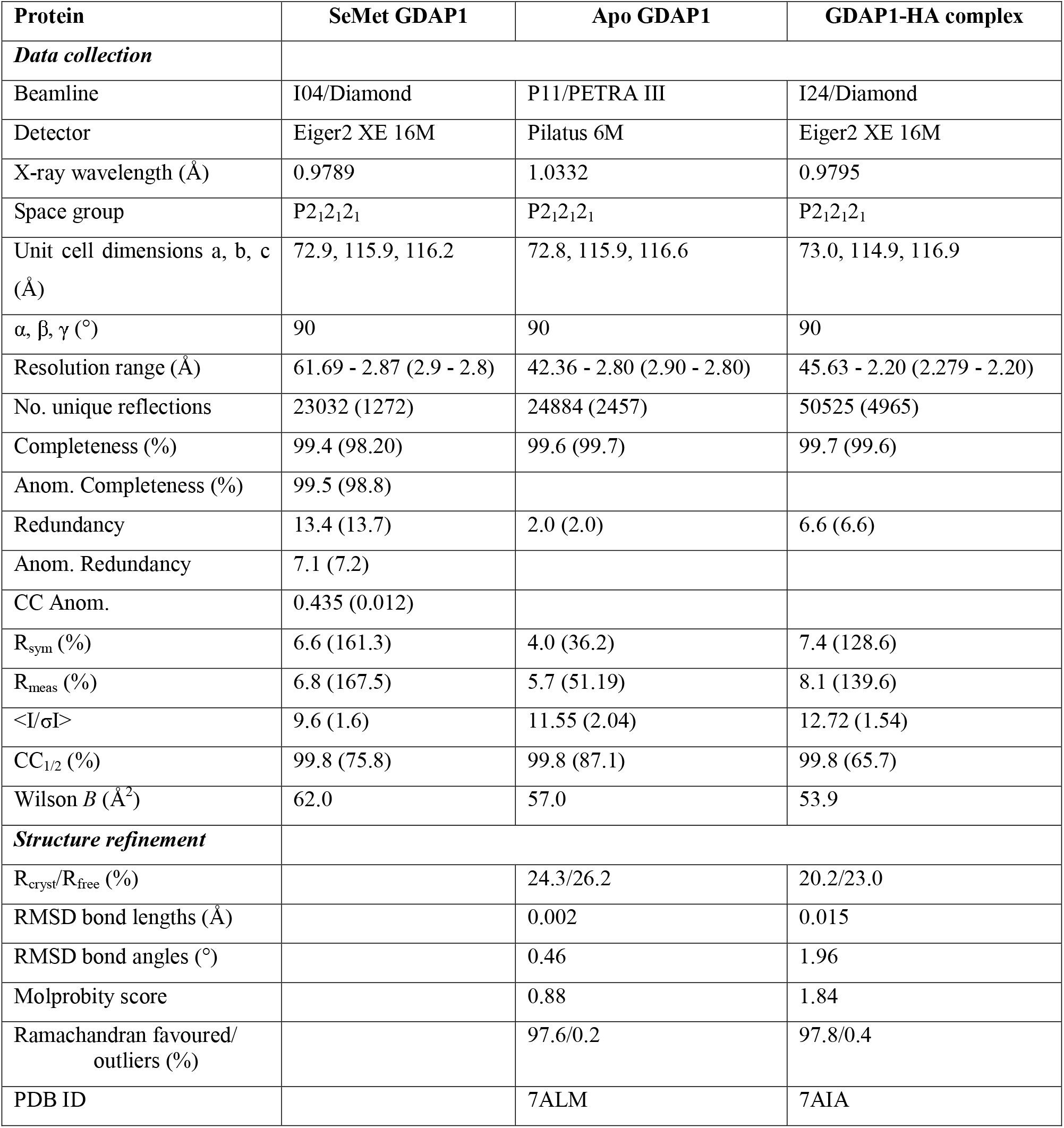
Diffraction data and refinement statistics.

**Figure 5.**
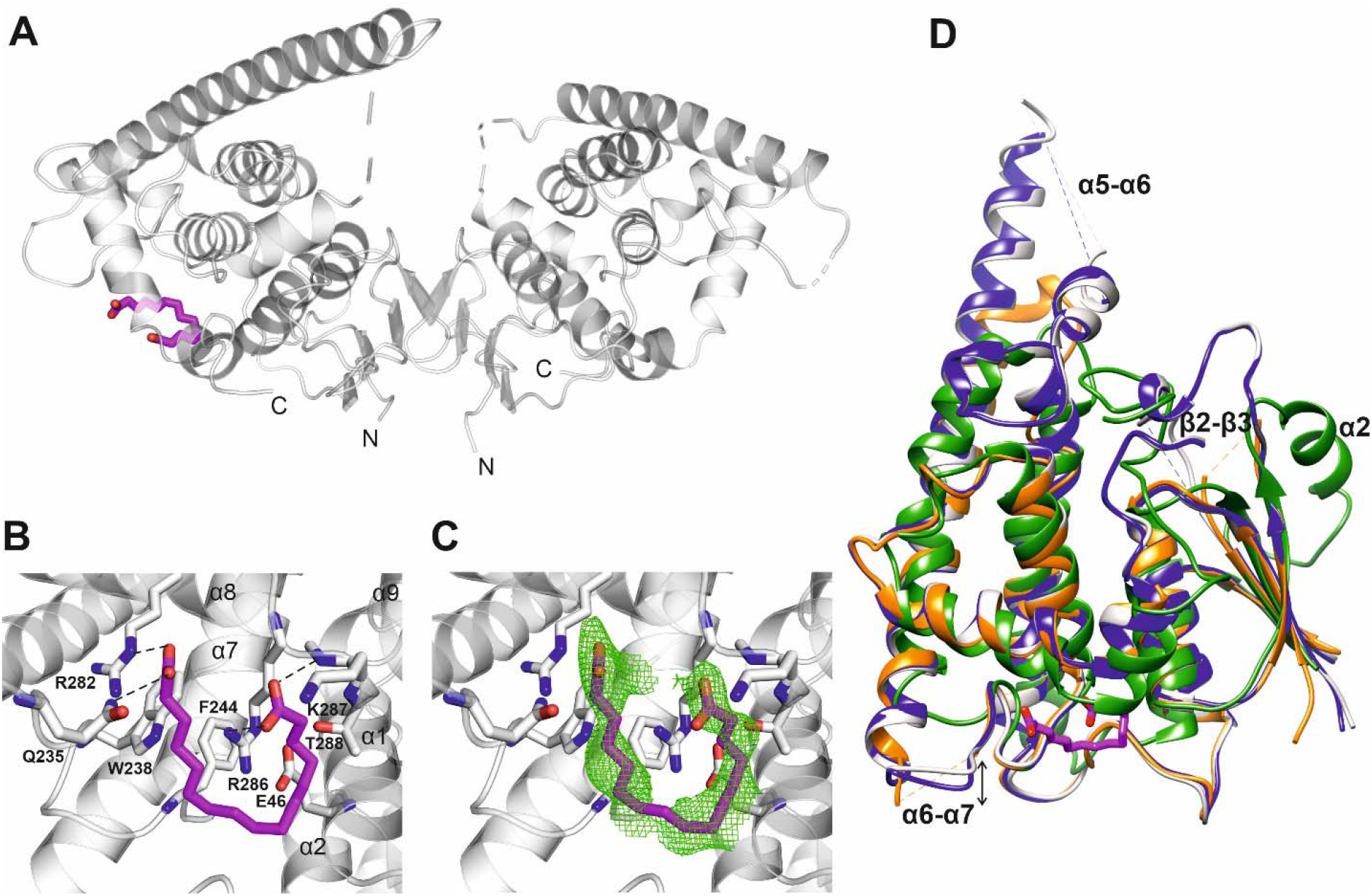
Crystal structure of GDAP1Δ303-358 in complex with HA. A. Overall structure of the GDAP1-HA complex. The dashed line indicates a loop not resolved in electron density. HA is shown in magenta. B. HA interacting residues, hydrogen bonds are shown as dashed lines. C. HA ligand of GDAP1 overlaid with Polder map (5σ, green). D. Superposition of apo GDAP1 (blue), the HA complex (grey), GDAP1 PDB ID 6uih ^7^ (orange) and sjGST PDB ID 1ua5 ^60^ (green). HA is shown in magenta. The dashed lines indicate loops not resolved in electron density. The mobile loops include β2-β3, α5-α6, and α6-α7.

To summarize, although the GDAP1 core domain and canonical GSTs share a similar monomer fold, the crystal structure of GDAP1 reveals a novel dimer interface. The lack of GST activity and GSH binding confirm that GDAP1 has a unique structure and function compared to the rest of the GST family. HA plays a role as an allosteric modulator of oligomerization, flexibility, and stability of GDAP1, at least *in vitro*.

### Identification of key residues for GDAP1 dimerization

The crystal structure of GDAP1 reveals Cys88 and Tyr29 as central for dimer formation (Figure 4D). To confirm their essential role of at the dimer interface, mutations were generated, including C88A, Y29F, Y29F/C88A, and Y29E/C88A, and oligomerization was investigated using SEC, SEC-MALS, and SEC-SAXS. Comparison of SEC elution profiles shows that the Y29F mutant retains a small amount of dimer, whereas the C88A and Y29F/C88A mutations significantly inhibit dimer formation (Figure 6A). In non-reducing SDS-PAGE, a dimer band is present for Y29F but absent for C88A and Y29F/C88A (Figure 6B). According to SEC-MALS, the main peak for wild-type GDAP1 is a dimer, whereas C88A GDAP1 is a mixture of dimer and monomer with an apparent mass of 55.3 ± 4.8 kDa, similar to the second peak of the wild-type protein (Figure 6C). To examine the stability of wild-type GDAP1 and the C88A mutant, we used nanoDSF, a label-free fluorimetric technique that can determine the thermostability of proteins by following changes in their intrinsic fluorescence. The T_m_ for wild-type GDAP1 and C88A were +62.1 ± 0.23 °C and +57.4 ± 0.01 °C, respectively, which probably reflects the larger dimer fraction of wild-type GDAP1. Taken together, Cys88 is important for GDAP1 dimerization and stability.

**Figure 6.**
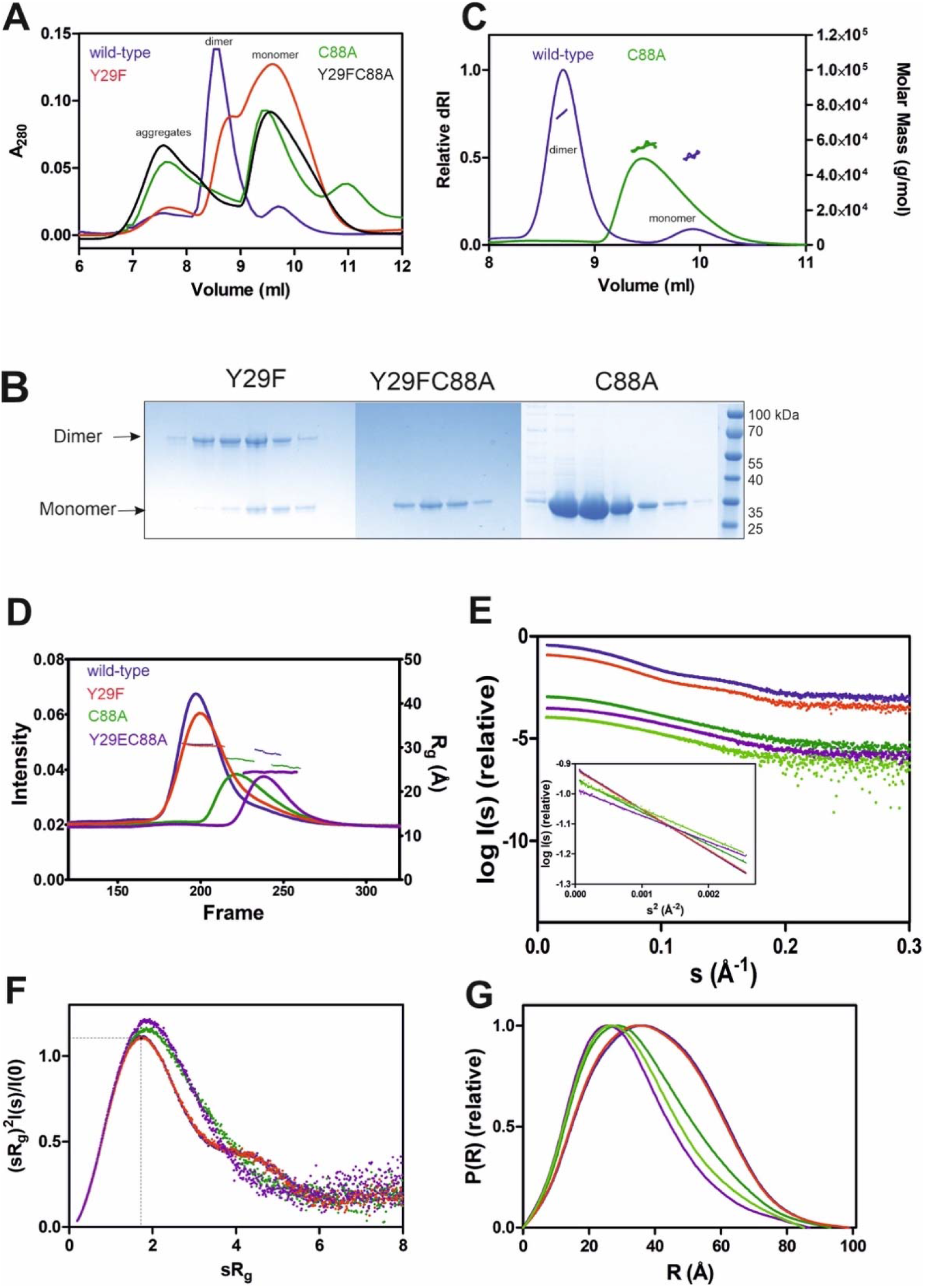
The oligomerization of wild-type and mutant GDAP1Δ303-358 using SEC, SEC-MALS and SEC-SAXS. A. SEC elution profile of wild-type (purple), C88A (green), Y29F (dark orange) and Y29FC88A (black) using column S75 increase 10/300 GL. B. Non-reducing SDS-PAGE gel of mutants SEC elution. C. SEC-MALS analysis of wild-type (purple) and C88A (green) using column S75 increase 10/300 GL. D. SEC-SAXS elution profiles and R_g_ plots. E. Experimental scattering data (log (I_s_) vs s) and linear fits in the Guinier regions (inset). The main and the second half of C88A peak are shown in dark and light green, respectively. F. R_g_ normalized Kratky plots, the dashed lines representing the maximum value of a standard globular protein G. Distance distributions *p(r)* plots of wild-type GDAP1Δ303-358 dimer (blue) and mutants Y29F (red), Y29EC88A (purple), C88A main peak (dark green) and the second half (light green).

To confirm the role of Tyr29 and Cys88 for GDAP1 dimerization, oligomerization of the mutants was studied using SEC-SAXS. Wild-type GDAP1 and the Y29F mutant eluted as a dimeric form, whereas C88A and Y29E/C88A eluted later (Figure 6D). In SAXS, Y29F at high concentration (10 mg/ml) shows a more substantial dimer peak in comparison to SEC data (Figure 6A – 5 mg/ml), implying that the hydrogen bond between Tyr29 residues is involved in dimerization, but not strictly required. The scattering curves and Guinier fits confirm sample monodispersity (Figure 6E).

According to its dimensionless Kratky plot, dimeric wild-type GDAP1Δ303-358 is compact, whereas the monomeric form observed for mutant proteins is less compact (Figure 6F). The D_max_ of the monomeric C88A and Y29E/C88A variants is shorter compared to dimeric wild-type GDAP1 (Figure 6G and Table 2). The main peak of C88A has a larger molecular weight compared to Y29E/C88A (Table 2), suggesting that this peak is a mixture of dimer and monomer, whereas the second part of the peak represents a monomer and shows a molecular weight and distance distribution similar to Y29E/C88A (Figure 6G, Table 2). The *ab initio* model of Y29E/C88A superimposes well with the crystal structure of GDAP1 chain A (Figure S9A). Particle shape reconstruction was done for the Y29E/C88A mutant (Fig S9B). According to its R_g_, it is monomeric. The *ab initio* map corresponds to a clearly non-spherical shape, indicating that the GDAP1 monomer exists in an extended conformation in solution.

For further insight into the structure of monomeric GDAP1, low-resolution electron density maps were reconstructed for the GDAP1Δ303-358 wild-type and the Y29E/C88A mutant. The averaged Y29E/C88A map reveals a monomeric particle, in line with the *ab initio* model (Figure S9B). The map reveals a shape similar to the monomeric mouse GDAP1 crystal structure 7.

Taken together, SEC, SEC-SAXS, and SEC-MALS data confirm that Cys88 plays an important role at the dimer interface. Tyr29 contributes with a regular hydrogen bond, a C-H…pi bond to Ile81, and a number of van der Waals interactions. The mutation Y29E/C88A abolishes the disulfide bond and disrupted the hydrophobic surface on the dimer interface, generating a monomeric form.

### GDAP1L1 is monomeric

GDAP1L1 is a paralogue of GDAP1 with 55% sequence identity (Figure S10) and is mainly cytosolic 15. As opposed to full-length GDAP1 (data not shown), full-length GDAP1L1 over-expressed in *E. coli* can be purified to homogeneity and is soluble (Figure 7A). Under both non-reducing and reducing conditions, GDAP1L1 migrates as a 44-kDa monomer on SDS-PAGE (Figure 7A). Mass spectrometry confirmed the protein band to be full-length GDAP1L1. Sequence alignments show that Cys88 and Glu84, involved in the dimer interface of GDAP1, are replaced by Ser109 and Asp105, respectively, in GDAP1L1 (Figure S10). Coupled with the high sequence similarity, GDAP1L1 folds like GDAP1 but does not for dimers.

**Figure 7.**
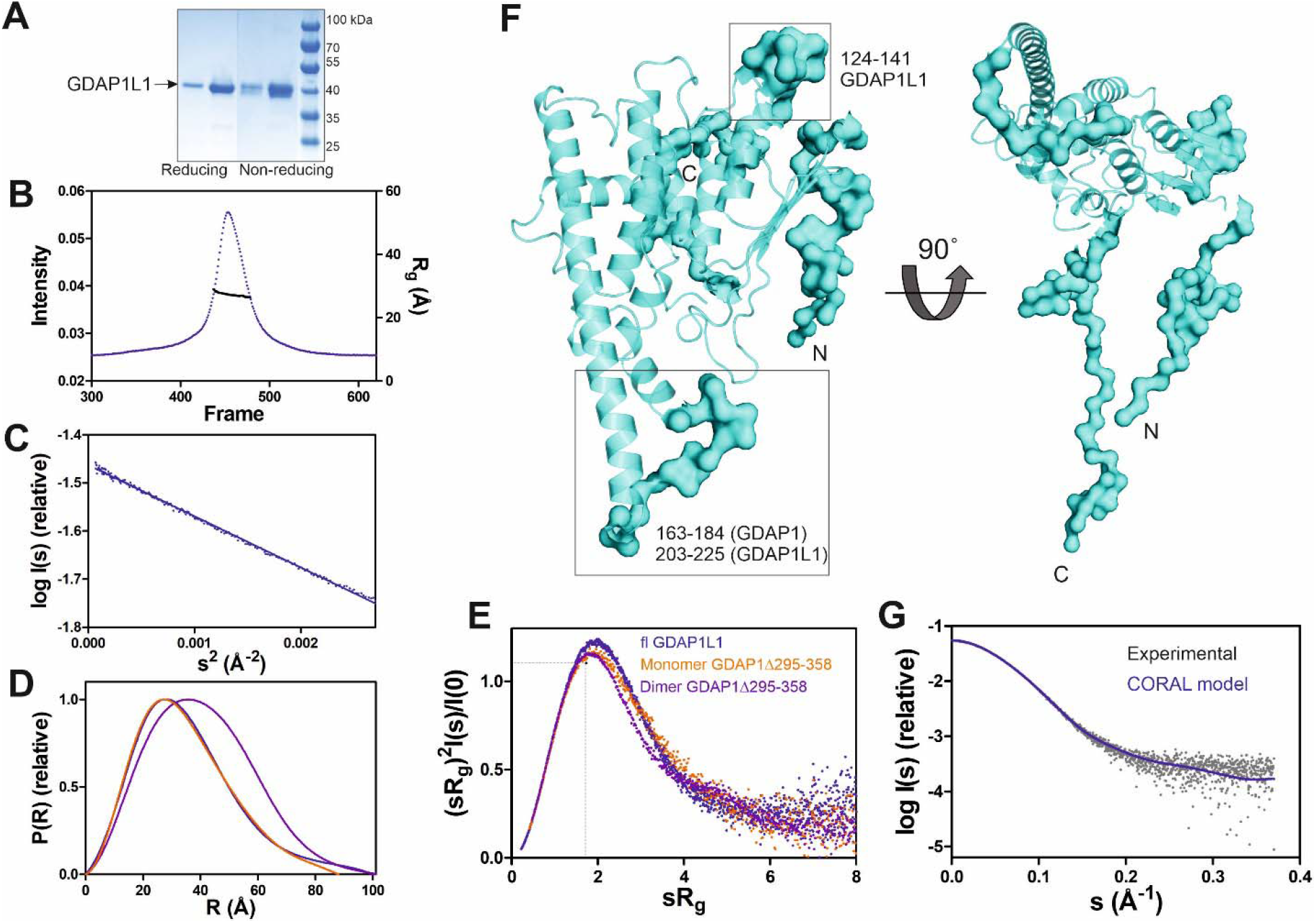
SAXS analysis of GDAP1L1. A. Non reducing and reducing SDS-PAGE gel of GDAP1L1. B. SEC-SAXS elution profile and R_g_ plot of SAXS frames. C. Guinier analysis. D. Distance distributions *p(r)* plots. E. Dimensionless Kratky plots for GDAP1L1 (blue) and GDAP1 monomer (orange) and dimer (purple). The dashed lines represent the peak position for a standard globular protein. F. CORAL model of GDAP1L1 based on GDAP1 crystal structure (cartoon), surface for the restored missing fragments. G. Experimental scattering curve of GDAP1L1 (grey) overlaid with the CORAL model (blue, χ^2^ =1.19) using CRYSOL.

SEC-SAXS was used to study the oligomeric state of GDAP1L1, revealing an R_g_ of 27 Å (Figure 7B). In line with this, the GDAP1L1 molecular mass calculated from volume correlation is 43.7 kDa, and from Bayesian estimate 44.7 kDa. A linear Guinier fit indicates that GDAP1L1 is quantitatively monomeric (Figure 7C). The P(r) function of GDAP1L1 has a similar shape as monomeric GDAP1Δ295-358, except for a long tail, leading to a D_max_ of 100 Å (Figure 7D). This tail implies that GDAP1L1 has disordered regions, most likely corresponding to the N terminus and the C-terminal HD and TMD. It thus seems that the single transmembrane domain does not make recombinant GDAP1L1 insoluble; this behaviour is different from GDAP1 and could be related to the different oligomeric state. The dimensionless Kratky plot of GDAP1L1 shows an asymmetric bell-shaped curve (Figure 7E), indicating increased structural flexibility compared to the GDAP1 core domain. A SAXS-based hybrid model of full-length GDAP1L1 was generated based on the GDAP1 crystal structure and complemented with the missing loops and termini (Figure 7F). This hybrid model fits the experimental data (Figure 7G) and shows flexible regions in addition to the folded monomeric core domain (Figure 7F).

## DISCUSSION

We carried out a detailed structural characterization of human GDAP1 containing the full GDAP1-specific insertion, containing the α5-α6 loop and the long α6 helix. The results indicate that GDAP1 forms a unique type of homodimer mediated by a hydrophobic surface and a disulphide bridge. Furthermore, a fatty acid ligand for GDAP1 was identified. Together with earlier data, our results provide important clues towards the structure and function of GDAP1 on the outer mitochondrial membrane and its involvement in neurodegenerative disease.

### GDAP1 is a unique member of the GST family

Although the sequence identity is only ~20%, the GDAP1 core domain shares a fold similar to canonical GST enzymes. However, GDAP1 has a unique mode of dimerization, and it lacks GST activity. The main differences constitute a missing α helix between β2 and β3 and the unique helices α5 and α6 with the connecting α5-α6 loop (Figure 4, 5, S6). Variations in these regions prevent GDAP1 from forming canonical GST dimers and interacting with typical GST substrates. Dimerization is critical for GST activity in all eight known GST classes ^61, 62^. Mutations at the GST dimer interface result in a stable, soluble, but inactive enzyme ^63^. The unique arrangement of the GDAP1 interface suggests a different function for GDAP1.

In α, μ, π, and *S. japonicum* GST, a “lock-and key” kind of hydrophobic interaction is established by wedging a hydrophobic side chain (Phe52, α; Phe56, μ; Phe47, π; Phe51, *S. japonicum*) from one monomer into a hydrophobic pocket on the second one, formed by five conserved residues on helices α4 and α5 64. In GDAP1, the “key” Phe and “lock” residues are not conserved (Figure S12).

The regions β2-α2-β3 and α4-α5 form the GSH binding site of GSTs, involving many interacting residues ^65^, which are not conserved in GDAP1 (Figure S11A, S12). Googins *et al.* identified differences between the G-sites of GDAP1 and canonical GSTs, including limited sequence conservation in the α2 region ^7^. Contrary to predictions, we show that GDAP1 lacks helix α2. Apo and GSH-bound GSTA1-1 show a conformation of the α2 helix, which is completely different from GDAP1 loop β2-β3 (Figure S11A). On the other hand, the catalytic Tyr9 residue of GSTA1-1 is conserved as Tyr29 in GDAP1, but Tyr29 points to another direction and makes central contacts at the GDAP1 dimer interface (Figure S11A). Hence, a similar fold makes GDAP1 a member of the GST enzyme family, but differences in the dimer interface and important residues for GSH binding and catalysis imply a unique function within the family.

### GDAP1 as a target for CMT mutations

A large number of CMT-related mutations in *GDAP1* have been identified. The most common *GDAP1* genotype in 99 Spanish patients was p.R120W ^66^. R120W, H123R, A156G, and P274L were reported in European patients ^67^. Several mutations have been studied using neurons and Schwann cells or a yeast model ^67–69^. The GDAP1 crystal structure now allows establishing a molecular basis for many of the known mutations in the human gene mutation database (http://www.hgmd.cf.ac.uk/ac). A CMT-related mutation cluster of GDAP1 mainly localizes on helices α3 and α6, and less on helices α7, α8, and their connecting loops (Figure 8). There are 46 published missense mutations involving 39 residues. The main cluster contains 27 residues that interact closely with each other in the crystal structure, including salt bridges, hydrogen bonds, and van der Waals interactions, forming a network of interactions (Figure 8). CMT mutations hindering these interactions could affect GDAP1 folding and stability, in addition to its interactions with other molecules. Interestingly, HA binds to GDAP1 in a pocket next to this cluster and forms a hydrogen bond with Arg282 (Figure S11B). HA binding increases GDAP1 stability by inducing a conformational change of the loop α6-α7, which is involved in the mutation cluster. Thus, the CMT-related cluster and HA binding site may relate to the function and/or folding of GDAP1. GDAP1 is highly conserved between vertebrates but not fruit fly (Figure S10). Crucial residues on the GDAP1 dimer interface, including Tyr29 and Cys88, and many CMT-related residues are conserved, suggesting a role in the structure and function of GDAP1. Further studies are needed to investigate GDAP1 function and its relation to CMT, and current structural data provide a strong basis for targeted experiments.

**Figure 8.**
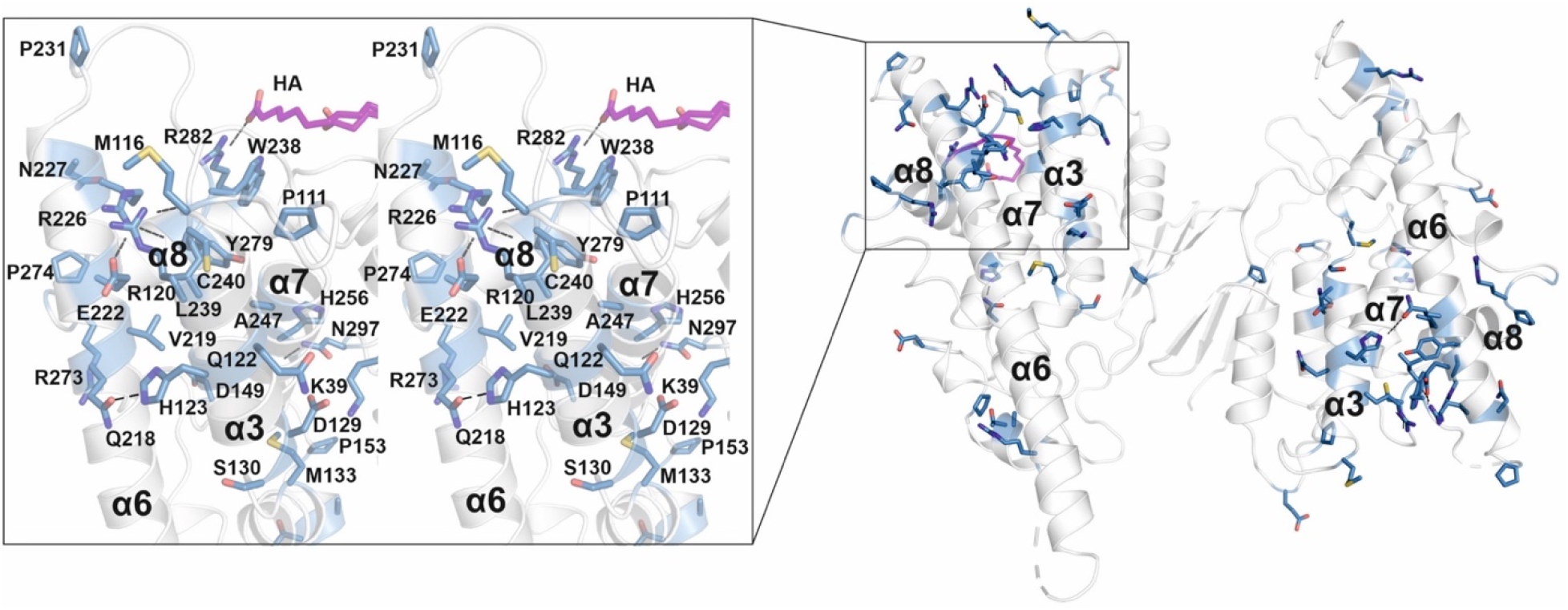
CMT-related residue cluster. Close-up stereo view of the CMT-related residue cluster (sticks) and HA binding site. Hydrogen bonds are shown in dashed line. HA is shown in magenta.

### GDAP1 and GDAP1L1 comparison

As a paralogue of GDAP1, GDAP1L1 shares a high sequence identity and the same fold. However, GDAP1L1 is monomeric and soluble (Figure 7). GDAP1 and GDAP1L1 have many conserved residues at the GDAP1 dimer interface, except for the central residues Cys88 and Glu84 (Figure S10). Gly83, a residue localized at the hydrophobic surface on the dimer interface, is replaced by Arg in GDAP1L1 (Figure S10). G83R is a CMT-related mutation in an Italian family ^70^. GDAP1L1 might have one additional α helix between α2 and α3 (Figure S10). A shorter C terminus could be linked to the observed solubility of full-length GDAP1L1 compared to GDAP1.

Due to the conserved HD and TMD (Figure S10), GDAP1L1 can target to mitochondria and compensate for GDAP1 deficiency ^14^. Hence, it appears that the HD and TMD are essential for GDAP1/GDAP1L1 mitochondrial targeting, while the GST-N and GST-C domains play a role in another function.

### Functional considerations

The unique α-loop of GDAP1 is involved in interactions with β-tubulin ^68, 71^, indicating that GDAP1 may participate in the interaction between mitochondria and microtubules. The CMT-related cluster and the HA binding site highlight an important region of GDAP1. This region could be a binding pocket for a substrate and co-factor to catalyze a reaction if the protein functions as an enzyme. The region could also be a contact surface with other proteins, such as β-tubulin. It has been shown that the interactions between GDAP1 and β-tubulin were highly increased for the GDAP1 mutants at the CMT-related cluster and the long α6, including R120Q, R120W, T157P, R161H, and R282C, pointing towards a gain-of-function mechanism that affects spindle formation ^68^. It was speculated that *via* interaction with GDAP1 and other fission proteins, microtubules could be important for the interaction between mitochondria and the cytoskeleton ^68^.

GDAP1 was reported to interact with Rab6B, a protein localized to the Golgi apparatus and distributed in Golgi and ER membranes ^72^, and with caytaxin, a protein involved in mitochondrial transport ^71^. The interaction between these proteins may be important for the localization of mitochondria close to SOCE sites ^73^. GDAP1 mutations in the α-loop could perturb the protein interactions, thus inhibits SOCE activity or stimulate abnormal ^73^.

GDAP1 is not only located in mitochondria, but also in mitochondria-associated membranes (MAMs), and it may play a role at the interface between mitochondria and the ER ^71^. GDAP1-linked CMT may be associated with abnormal distribution and movement of mitochondria along the cytoskeleton towards the ER and subplasmalemmal microdomains ^71^. The bidirectional movement of lipids between the ER and mitochondria may be mediated by interactions between MAM and mitochondria ^74^. Fatty acids are a source of metabolic energy and function as building blocks for complex lipids. GDAP1 could be a fatty acid transport protein due to its localization on MAM and MOM and its fatty acid binding shown here. Moreover, since both GDAP1 and HA are linked to Ca^2+^ homeostasis, GDAP1 may regulate this metabolism through its binding to fatty acids.

Another aspect arising from these findings is the oligomeric state of GDAP1 *in vivo*. As shown by earlier studies from neuronal cell line protein extracts ^5^, GDAP1 seems to be expressed as a dimer, and our results show that the dimers are covalently bonded. Changes in the redox environment could easily alter this equilibrium. In cells, mutant variants of GDAP1 lead to depleted GSH levels, causing excess reactive oxygen species (ROS) stress, suggesting that GDAP1 may actively regulate GSH metabolism. These could affect mitochondrial membrane integrity and oxidative phosphorylation efficiency *via* an unknown mechanism ^9, 75, 76^. Thus, the oligomerization of GDAP1 could be a regulated event induced by specific ROS-sensitive pathways.

### Insights into the structure of full-length GDAP1

The crystal structure of the dimeric GDAP1 core domain lacks the HD and TMD, but full-length GDAP1 does form dimers in cells. Co-immunoprecipitation of full-length GDAP1 from HEK-293T cells confirmed that the protein formed homodimers ^6^. We built a model of dimeric full-length GDAP1 on a phospholipid membrane using the crystal structure of the complete human GDAP1 core domain (Figure 9). The transmembrane domain of GDAP1 contains a Gly zipper, a motif linked to the dimerization of transmembrane helices ^77^. The lipid fraction of MOM in mammals consists mainly of phosphatidylcholine, phosphatidylethanolamine, and phosphatidylinositol ^78^, with minor amounts of phosphatidylserine, cardiolipin, and phosphatidic acid. On both sides of the membrane, the model shows positively charged surfaces of the protein at the bilayer headgroup regions. The GDAP1-specific insertion has a strong positive potential and could be involved in molecular interactions, for example with the cytoskeleton.

**Figure 9.**
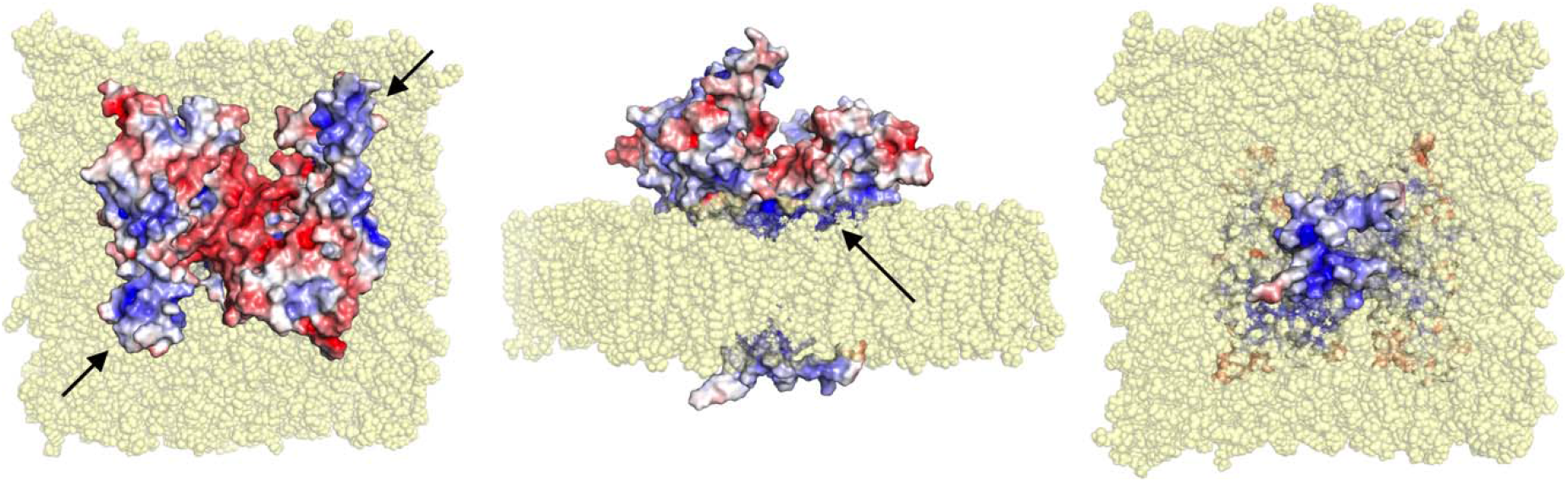
Model for full-length dimeric GDAP1 on a membrane bilayer. Full-length GDAP1, including modelled TMD and loops missing from the crystal structure is shown as an electrostatic potential surface. In the top view (left), note how the cavity at the dimer interface has negative potential, while the long helices and the ⍰5-⍰6 loop (arrows) have positive charge. The side view (middle) shows a cavity in the dimer arrangement towards the cytosol (top) and a positively charged surface facing the membrane (arrow). The bottom view (right) indicates that the tail is very small and positively charged.

The fatty acid binding site observed in the GDAP1 crystal structure faces the membrane-binding surface in the context of the modeled full-length dimer, suggesting that the observed ligand could mimic the lipid membrane surface. This in turn suggests that membrane binding could affect the conformation of the GDAP1 core domain, for example, *via* the incorporation of acidic lipid headgroups in the binding site. These questions can be answered when the structure of full-length GDAP1 on a MOM-like lipid membrane eventually becomes available.

### Concluding remarks

GDAP1 is linked to CMT and a member of the GST family; however, its function remains unclear at the molecular level. The crystal structure of the complete human GDAP1 core domain reveals a GST fold, with a previously unseen mode for dimerization. The monomer-dimer equilibrium could be further linked to redox phenomena in the cell, and the function of full-length GDAP1 on the MOM may be regulated by the oligomeric state. The GDAP1 structure and the discovery of the first GDAP1 ligand not only provide information to map the CMT-related residue cluster and the corresponding interactions in detail, but also provides a template conformation for further functional studies and structure-assisted ligand design. Further studies on GDAP1-linked CMT should use the human GDAP1 crystal structure as a reference framework to explain effects of mutations at the molecular level.

## Acknowledgements

This work was funded by the Academy of Finland, project number 24302881. We thank the core facilities at Biocenter Oulu, Finland, for technical support. Synchrotron beamtime and support at EMBL/DESY, Diamond Light Source, and SOLEIL are gratefully acknowledged. The research leading to this result has been supported by the project CALIPSOplus under the Grant Agreement 730872, as well as by iNEXT, grant number 653706, from the EU Framework Programme for Research and Innovation HORIZON 2020.

