## Supplementary Information for "Structure of the complete dimeric human GDAP1 core domain provides insights into ligand binding and clustering of disease mutations"

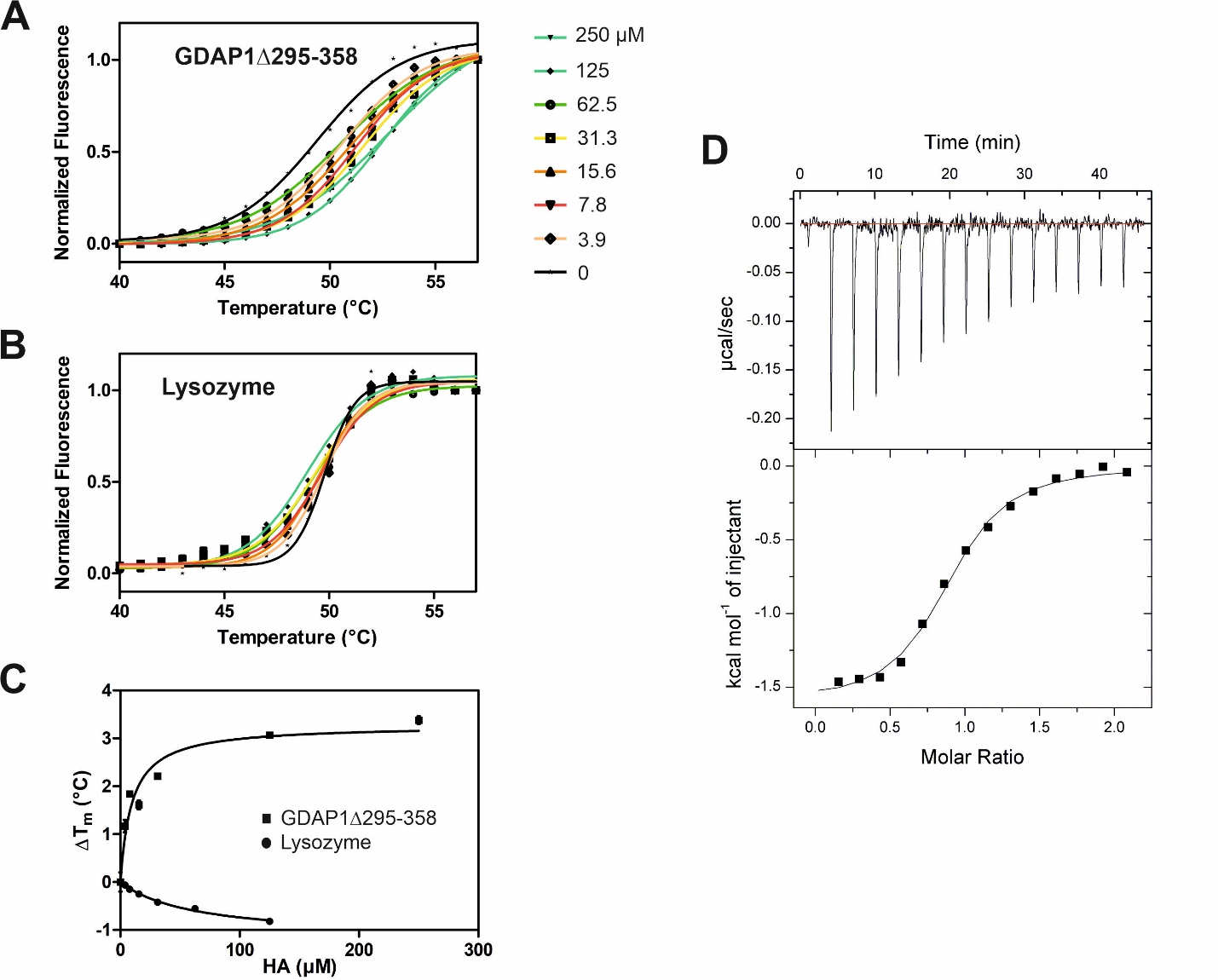

**Figure S1. HA binding stabilizes GDAP1∆295-358**

A. Thermal unfolding data of GDAP1∆295-358
B. Thermal unfolding data of lysozyme as a control
C. T_m_ shifts of GDAP1∆295-358 upon HA titration
D. ITC binding curve of HA binding to GDAP1∆295-358

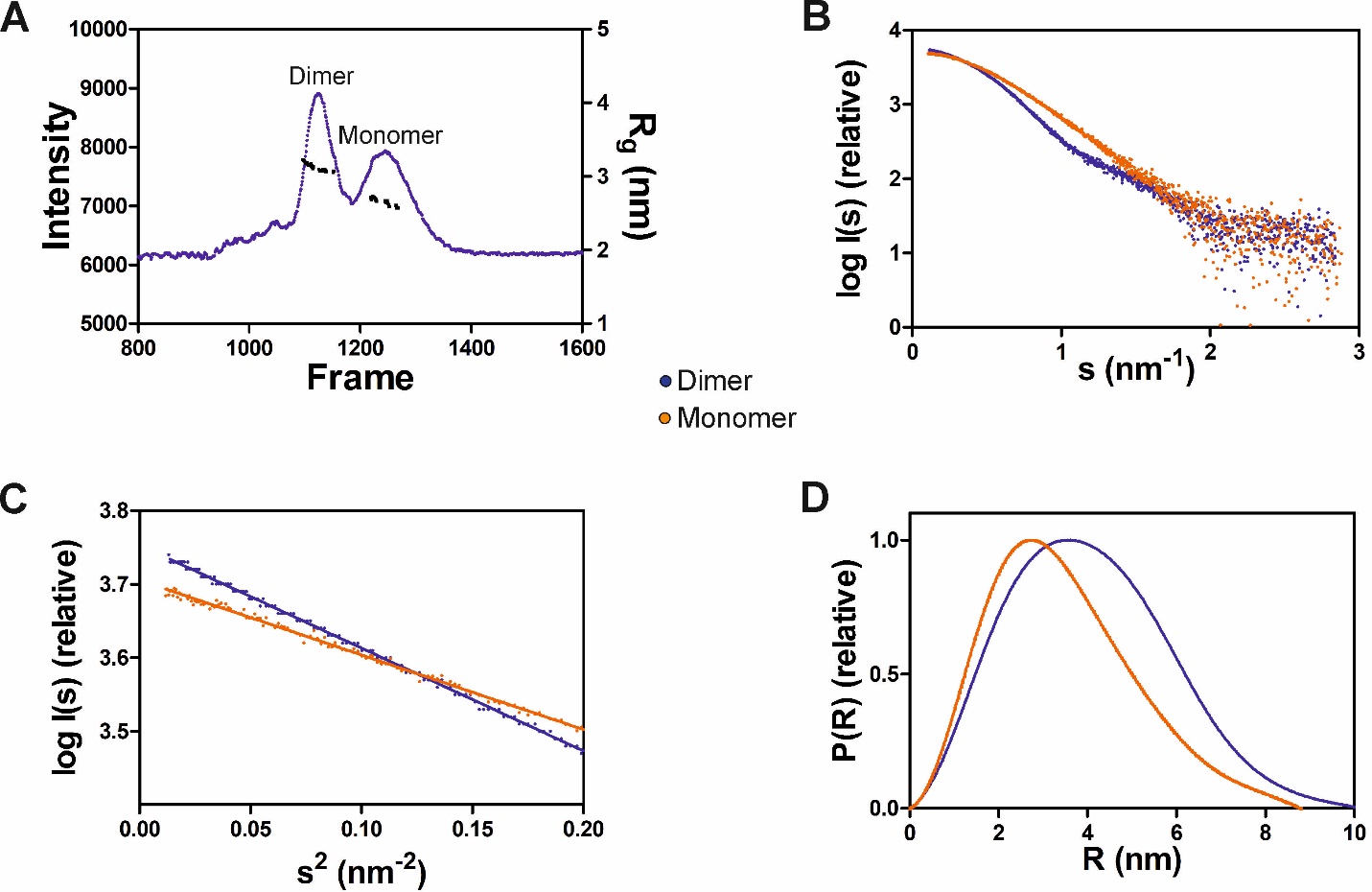

**Figure S2. SAXS analysis of GDAP1∆295-358**

A. SEC-SAXS elution profile. R_g_ for the dimer and monomer peaks is also plotted.
B. Experimental scattering data (log (I_s_) *vs*. s)
C. Guinier analysis
D. Distance distribution function for dimer (blue) and monomer (orange)

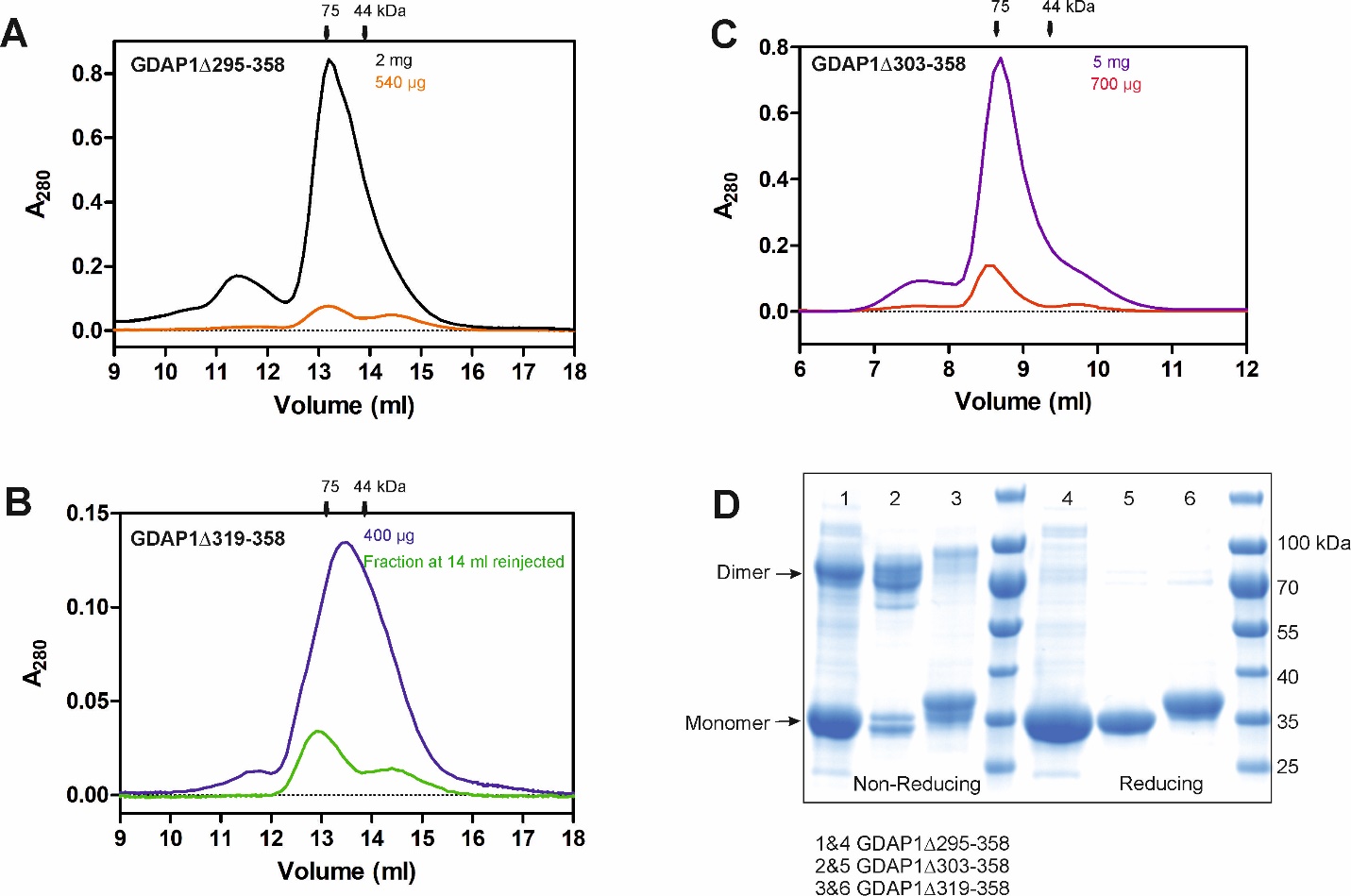

**Figure S3.** SEC elution profiles and SDS-PAGE gels. The positions of the molecular weight markers are shown on top.

A. Elution profile of GDAP1∆295-358 at 2 mg (black) and 540 µg (orange), column S200 increase 10/300 GL
B. Elution profile of GDAP1∆319-358 at 400 µg (blue) and ~100 µg (green), column S200 increase 10/300 GL
C. Elution profile of GDAP1∆303-358 at 5 mg (purple) and 700 µg (dark orange), column S75 increase 10/300 GL
D. Non-reducing and reducing SDS-PAGE gel of three constructs.

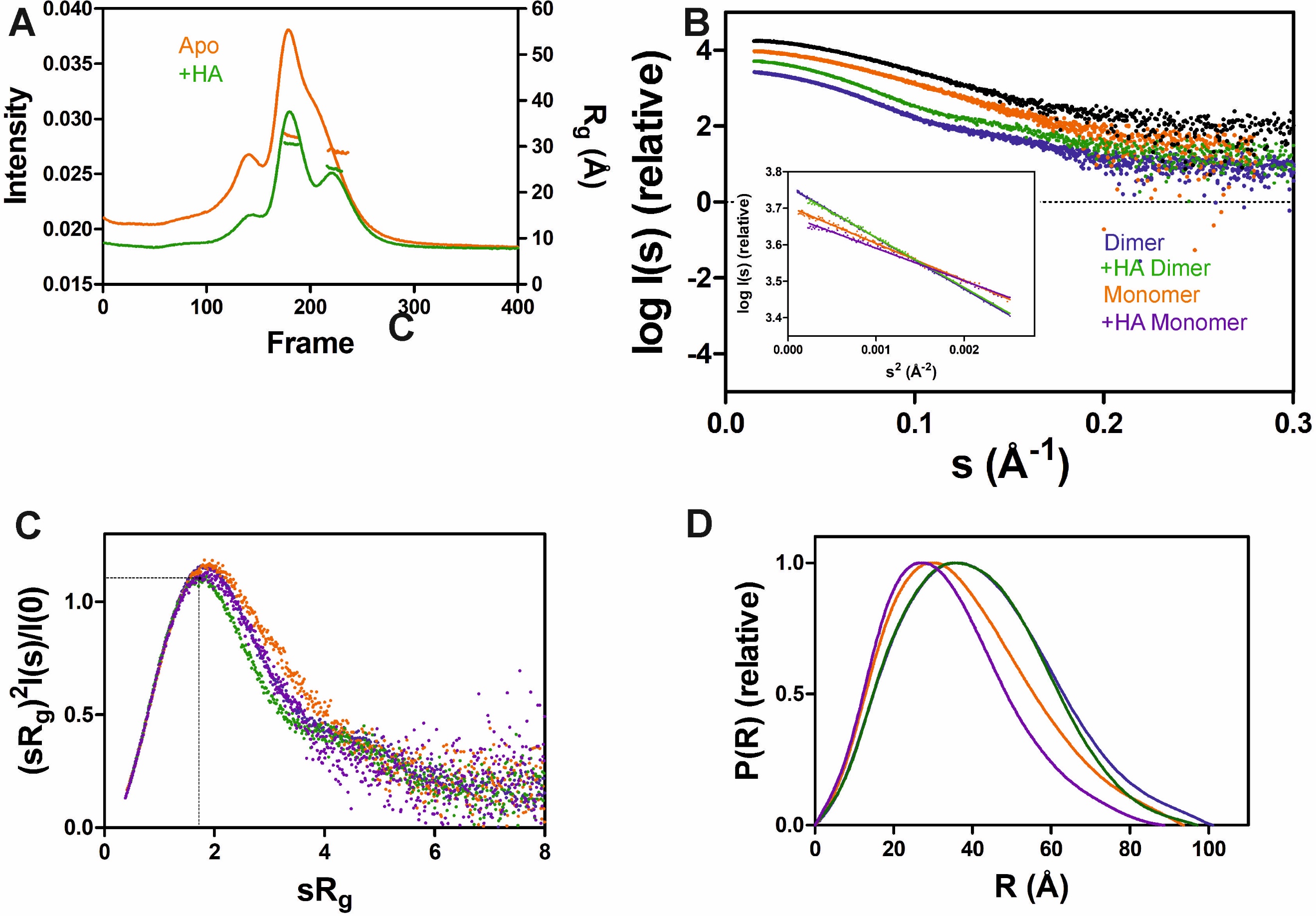

**Figure S4. SAXS analysis of GDAP1∆295-358 in the absence and presence of HA**

A. In-line SEC-SAXS elution profiles and R_g_ plot of SAXS frames for dimer and monomer peaks of the protein
B. Experimental scattering data (log (I_s_) vs s) and Guinier analysis (inset)
C. R_g_ normalized Kratky plots, the dashed lines representing the maximum value of the standard globular protein.
D. Distance distributions *p(r)* plots of ligand-free GDAP1 dimer (blue) and monomer (orange), ligand-bound GDAP1 dimer (green) and monomer (purple)

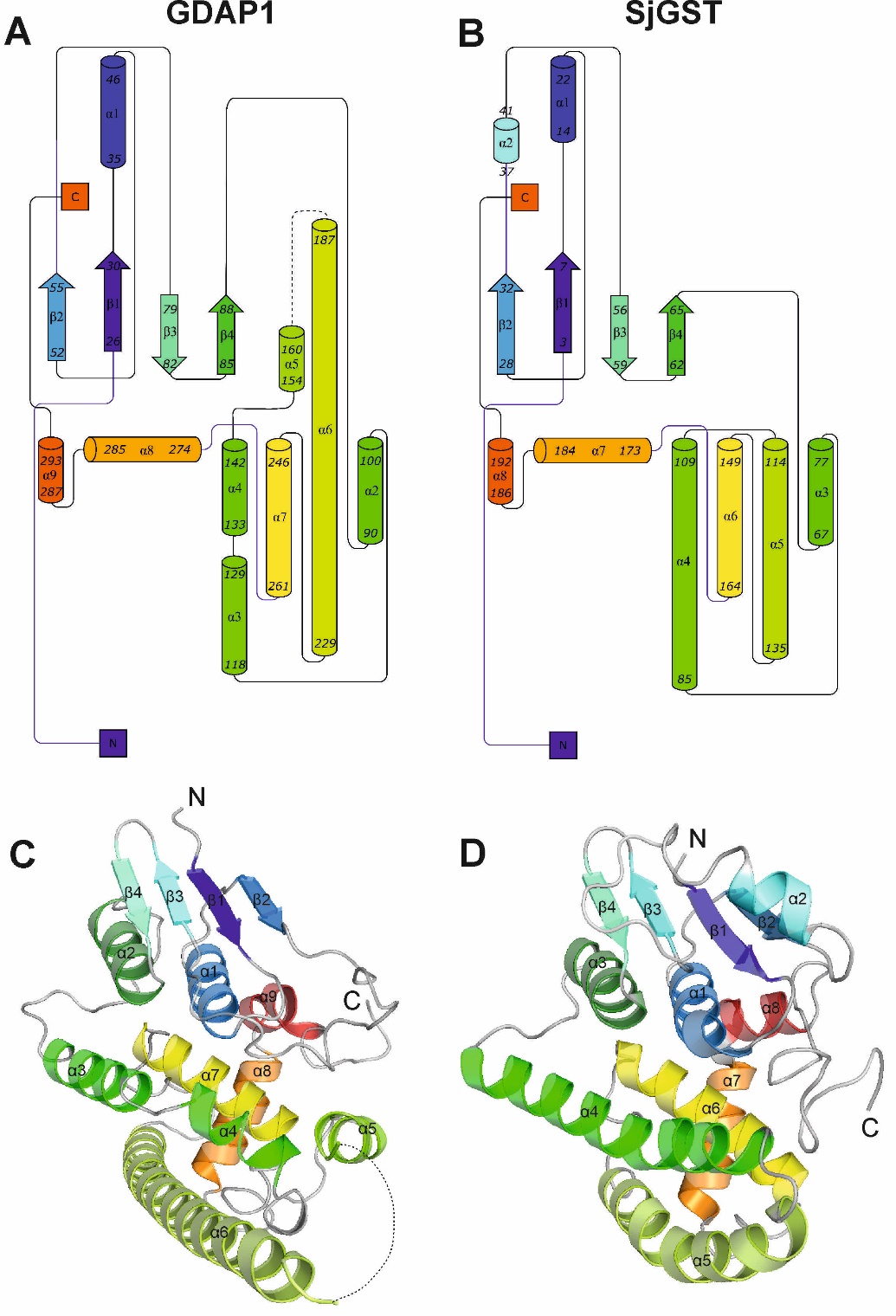

**Figure S5.** **Topology and structure of GDAP1 core domain and SjGST**

A, B. Topology diagrams for GDAP1∆303-358 chain A and SjGST (PDB:1UA5 ^1^). The dashed lines indicate loops not resolved in the electron density.

C, D. Cartoon representation of GDAP1 chain A and SjGST colored with gradient according to the topology diagrams. The dashed lines indicate loops not defined by electron density.

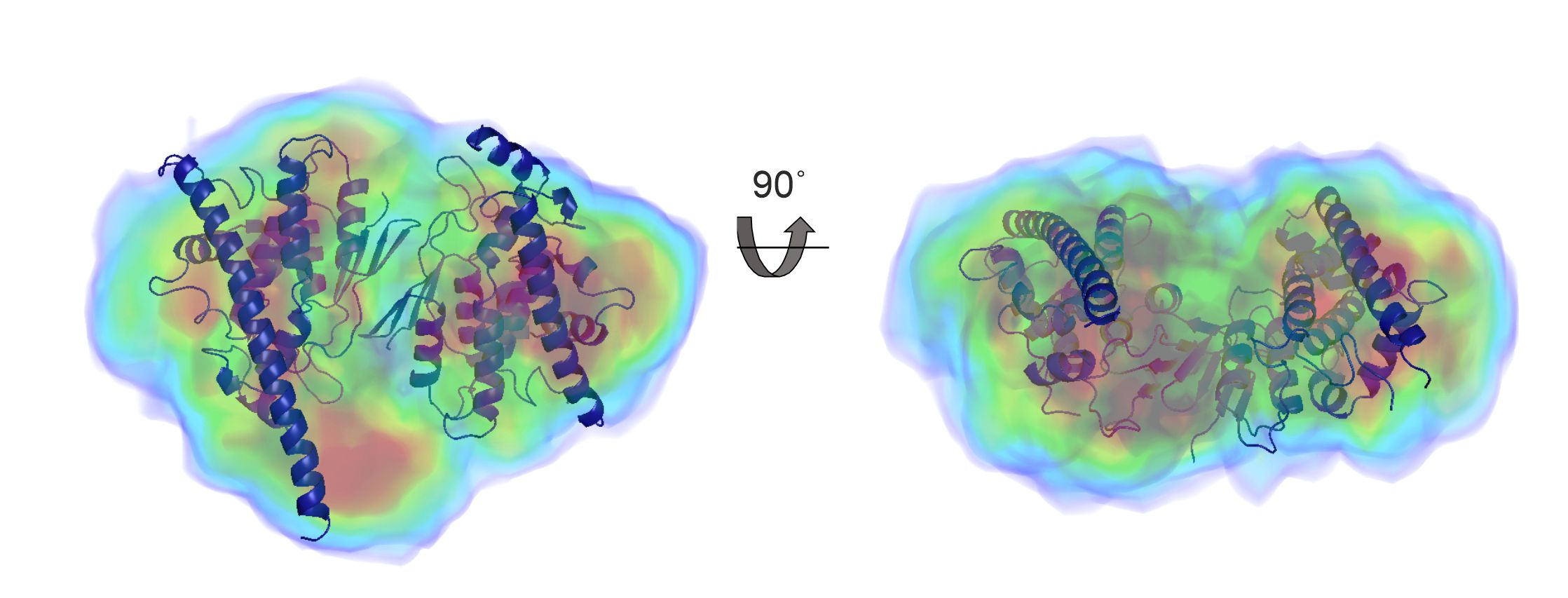

**Figure S6. Single particle electron density map reconstruction**

Single particle electron density map reconstruction (multicolor volume) and crystal structure alignment (blue) calculation using DENSS. The mean of 20 iterative map calculations were performed including enantiomer search. The median fit to the scattering data χ^2^= 1.455, and R_g_= 32.1 Å. The calculated mean support volume of the particle was 197080.59 Å^3^. The map resolution estimate was calculated with Fourier shell correlation function with a cut off value of 0.5. The mean resolution of 20 maps after refinement was 34.2 Å

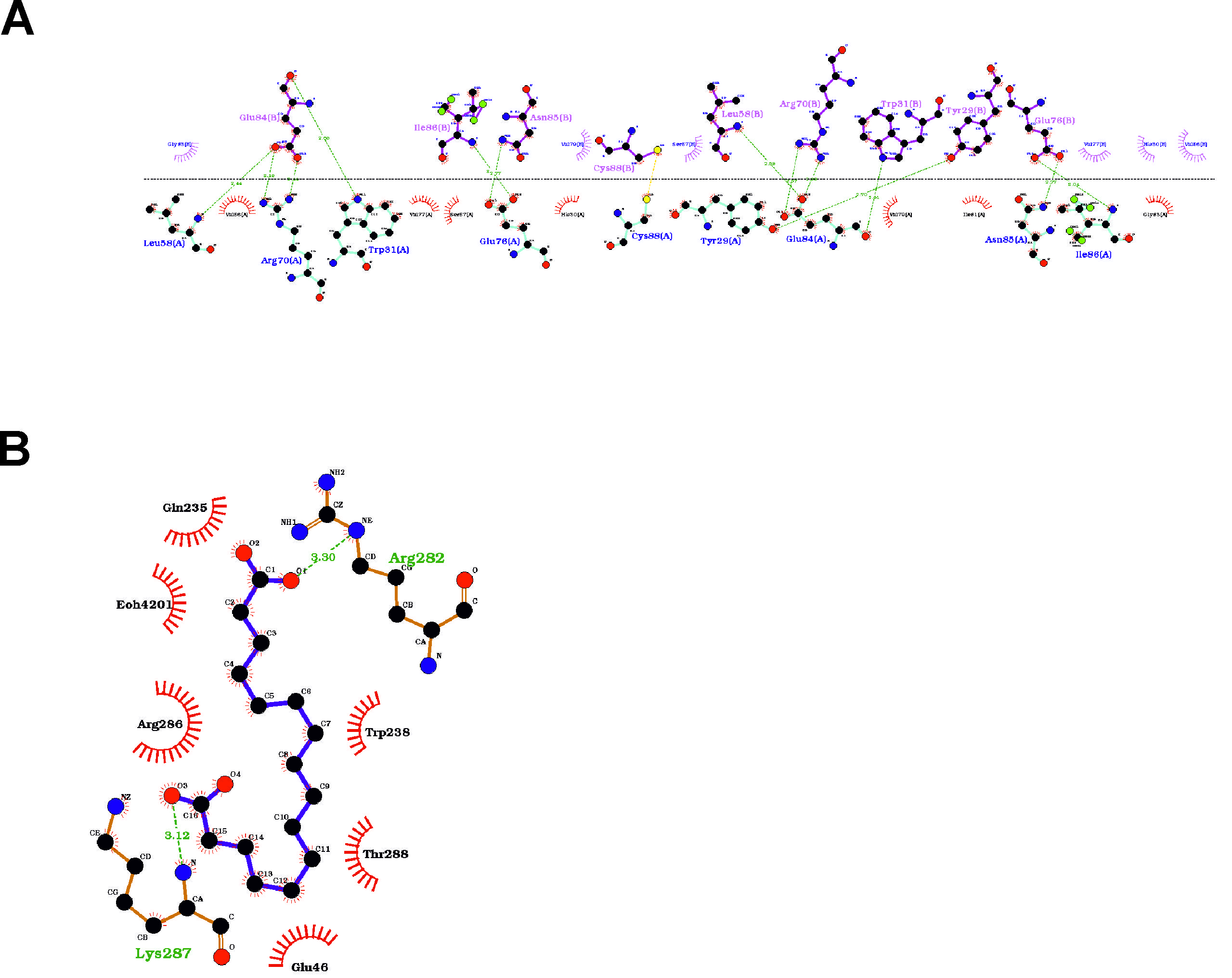

**Figure S7. Schematic view of interactions**

A. on the dimer interface
B. between HA and GDAP1

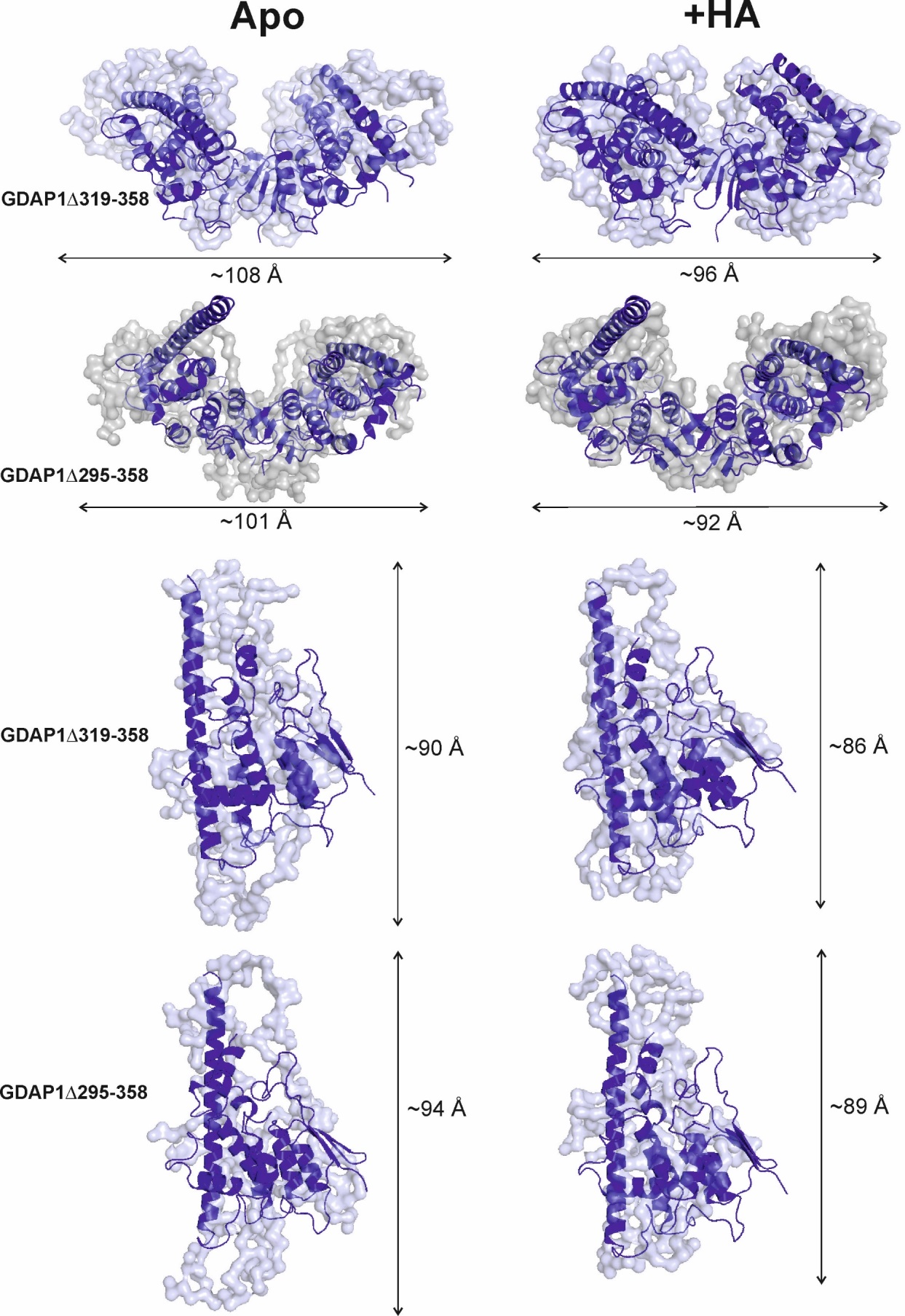

**Figure S8.** Chain-like models of GDAP1∆319-358 and GDAP1∆295-358 dimer and monomer in the absence and presence of HA with the respective maximum dimensions

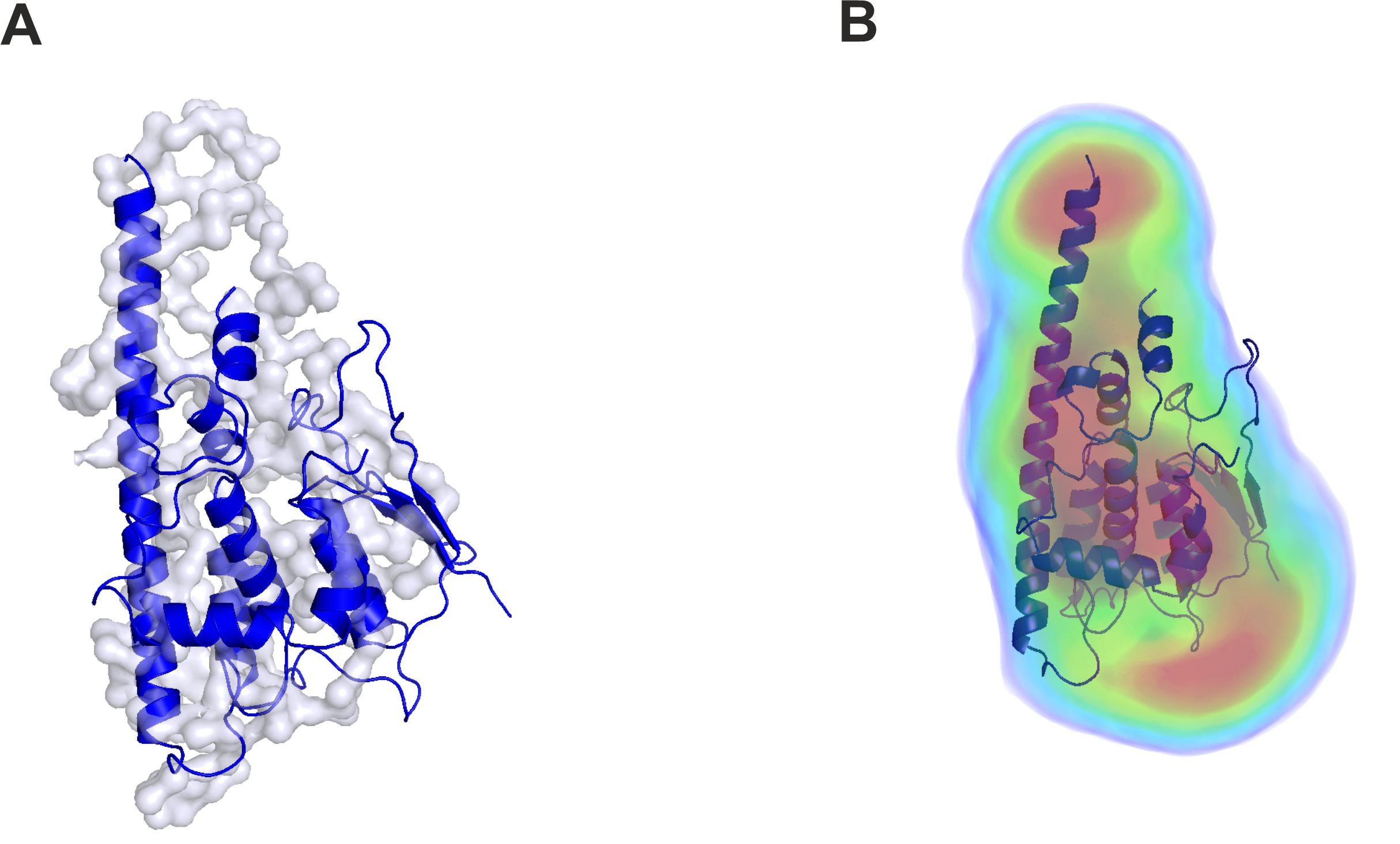

**Figure S9. *Ab initio* GASBOR models of mutant Y29E/C88A, and electron density map reconstruction calculated using DENSS.**

The Y29E/C88A GASBOR fit was χ^2^= 1.301, electron density map median fit χ^2^= 0.129, R_g_= 26.3 and the map resolution estimate 45.6 Å where FSC=0.5.

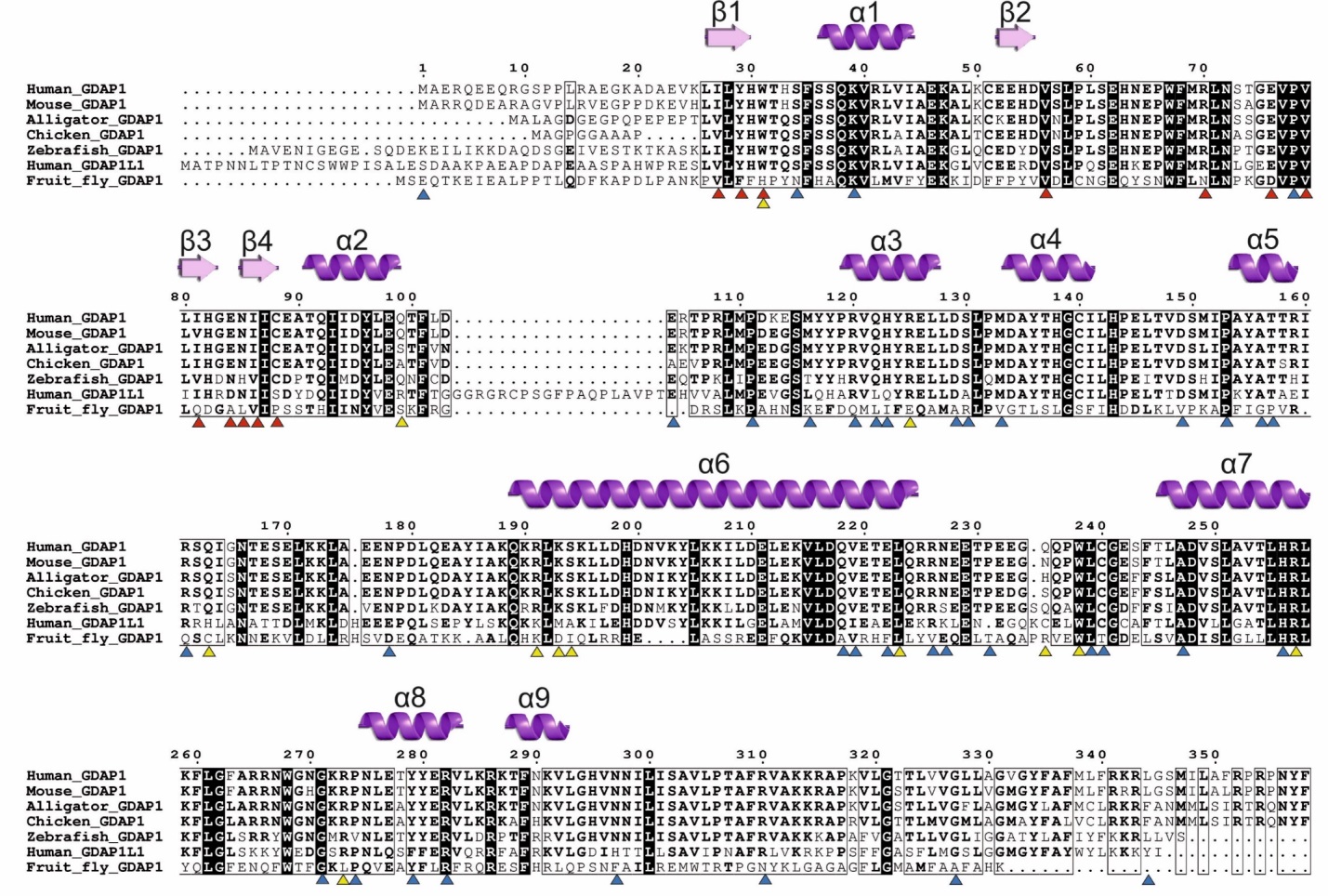

**Figure S10. Sequence alignment of GDAP1 and GDAP1L1**

Important residues for dimer interface is highlighted in red triangles and those implicated in CMT missense and nonsense mutations (based on the human gene mutation database <http://www.hgmd.cf.ac.uk/ac>) are highlighted using the blue and yellow triangles, respectively.

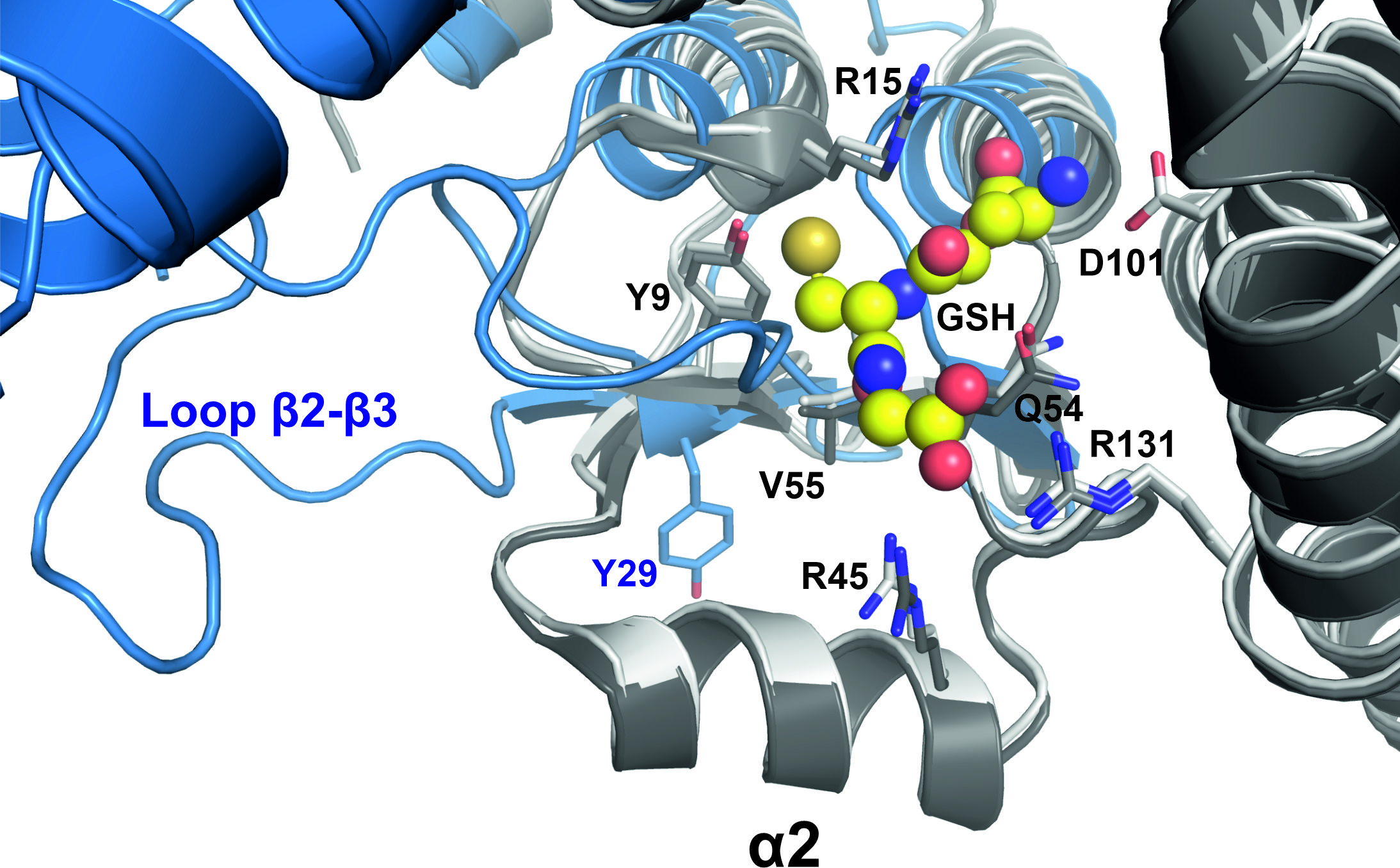

**Figure S11.** Superposition of GDAP1 with GSTA1-1

Close-up view of the GDAP1 loop β2-β3 (blue) and GSTA1-1 α2 of apo (dark grey, PDB ID 1pkz) and GSH-bound (light-grey, PDB ID 1pkw) ^2^. Key GST residues were shown in sticks and GSH was shown in spheres.

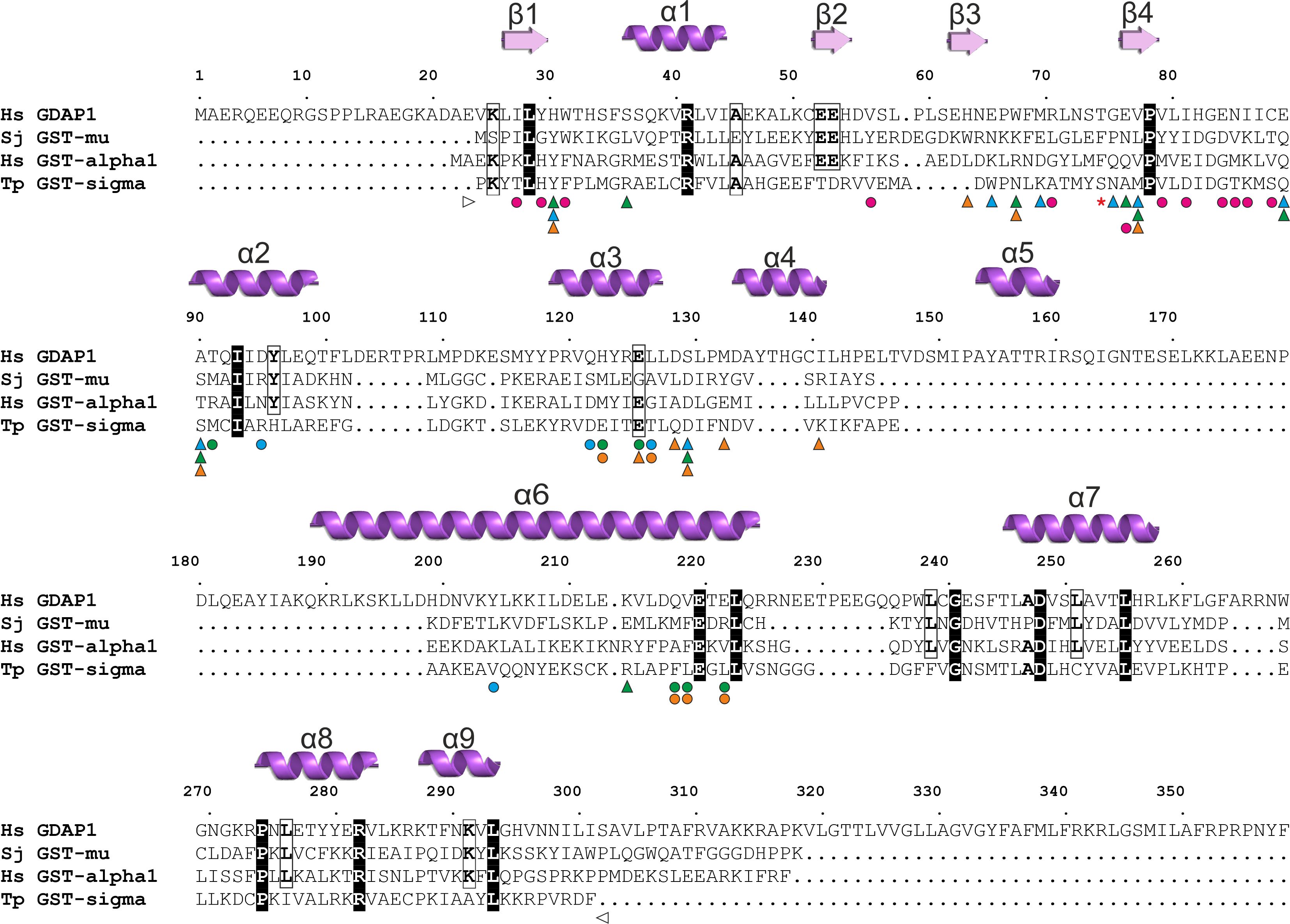

**Figure S12.** Sequence alignment of generic GSTs and GDAP1 (PDB ID: 7ALM, secondary structure elements). The alignment highlights key substrate interacting residues in GSTs (triangles) and dimer interface residues in GDAP1 and GSTs (circles). Additionally, the “lock-and-key”-type binding pocket residue present in GSTs is annotated with a red asterisk. The range of the GDAP1 crystal structure is marked with sided white triangles. Annotations of the GSTs are marked as *Schistosoma japonicum* GST-mu with orange (PDB ID: 1UA5), *Homo sapiens* GST-A1 with green (PDB ID: 1PKW), and *Todarodes pacificus* GST sigma with cyan (PDB ID: 1GSQ), respectively. The GDAP1 dimer interface residues are annotated as purple circles.

**Table S1. SASBDB entry codes**

| SASDJR8 | Dimeric human ganglioside-induced differentiation-associated protein 1, construct GDAP1∆295-358 |
| --- | --- |
| SASDJS8 | Dimeric human ganglioside-induced differentiation-associated protein 1, construct GDAP1∆295-358 with hexadecanedioic acid |
| SASDJT8 | Monomeric human ganglioside-induced differentiation-associated protein 1, construct GDAP1∆295-358 |
| SASDJU8 | Monomeric human ganglioside-induced differentiation-associated protein 1, construct GDAP1∆295-358 with hexadecanedioic acid |
| SASDJV8 | Dimeric human ganglioside-induced differentiation-associated protein 1, construct GDAP1∆303-358 |
| SASDJW8 | Monomeric human ganglioside-induced differentiation-associated protein 1, construct GDAP1∆303-358, mutant Y29E/C88A |
| SASDJX8 | Dimeric human ganglioside-induced differentiation-associated protein 1, construct GDAP1∆319-358 |
| SASDJY8 | Dimeric human ganglioside-induced differentiation-associated protein 1, construct GDAP1∆319-358 with hexadecanedioic acid |
| SASDJZ8 | Monomeric human ganglioside-induced differentiation-associated protein 1, construct GDAP1∆319-358 |
| SASDJ29 | Monomeric human ganglioside-induced differentiation-associated protein 1, construct GDAP1∆319-358 with hexadecanedioic acid |
| SASDJ39 | Monomeric human ganglioside-induced differentiation-associated protein 1-like 1, GDAP1L1 |

**Table S2. Electron density map reconstruction from SAXS scattering curves of wild-type and Y29EC88A mutant GDAP1∆303-358**

| **Structural parameters** | **WT** | **Y29EC88A** |
| --- | --- | --- |
| **q-range (s = 4π sin(θ)/λ)** | 0-0.504 | 0-0.327 |
| **2-fold symmetry** | Yes | No |
| **Dmax allowed (Å)** | 96 | 91 |
| **Real space Box width/range (Å)** | 228/±144 | 234/±117.5 |
| **Real space voxel volume (Å^3^)** | 23887872 | 12966281 |
| **Real space voxel size (Å)** | 4.5 | 3.6 |
| **Real space voxel volume (Å^3^)** | 91.1 | 49.5 |
| **Chi^2^** | 1.455 | 1.301 |
| **R_g_** | 32.1 | 26.3 |
| **Map resolution (FCS= 0.5)** | 34.2 | 45.6 |

**Table S3. GST activity measurement**

|  | **CDNB (340 nm)** | | | | **NBC (360 nm)** | | | | **EPNP (310 nm)** | |
| --- | --- | --- | --- | --- | --- | --- | --- | --- | --- | --- |
|  | **k2**  **(s^-1^)** | **±** | **Specific activity (nmol/min/µg)** | **±** | **k2**  **(sec-1)** | **±** | **Specific activity (nmol/min/µg)** | **±** | **k2**  **(sec-1)** | **Specific activity (nmol/min/µg)** |
| **GST** | 11.7 | 1.55 | 26.1 | 8.5 | 0.12 | 0.06 | 0.27 | 0.075 | n.d | n.d |
| **GDAP1∆295-358** | 7E-04 | 8E-04 | 2E-02 | 2E-02 | 6.005E-04 | 1.147E-04 | 1.334E-03 | 2.549E-04 | n.d | n.d |
| **GDAP1∆303-358** | 8.89E-03 | 1.25E-03 | 1.98E-02 | 2.77E-03 | 2.553E-03 | 1.686E-03 | 5.673E-03 | 3.747E-03 | n.d | n.d |
| **GDAP1∆319-358** | 3.81E-03 | 8.52E-05 | 6.35E-04 | 1.42E-05 | 1.631E-03 | 1.416E-03 | 3.625E-03 | 3.146E-03 | n.d | n.d |
